# Bayesian meta-analysis across genome-wide association studies of diverse phenotypes

**DOI:** 10.1101/477828

**Authors:** Holly Trochet, Matti Pirinen, Gavin Band, Luke Jostins, Wellcome Trust Case-Control Consortium 2, Gilean McVean, Chris C. A. Spencer

**Affiliations:** Wellcome Trust Centre for Human Genetics, University of Oxford, Oxford, UK; Institut de Cardiologie de Montréal (Centre de Recherche), Université de Montréal, Montréal, Québec, Canada; Institute for Molecular Medicine Finland (FIMM), University of Helsinki, Helsinki, Finland; Helsinki Institute for Information Technology HIIT and Department of Mathematics and Statistics, University of Helsinki, Helsinki, Finland; Department of Public Health, University of Helsinki, Helsinki, Finland; www.wtccc.org.uk; Big Data Institute, Li Ka Shing Centre for Health Information and Discovery, University of Oxford, Roosevelt Drive, Oxford, UK; Kennedy Institute of Rheumatology, University of Oxford, Roosevelt Drive, Oxford, UK; Christ Church, University of Oxford, St Aldates, Oxford, UK

## Abstract

Genome-wide association studies (GWAS) are a powerful tool for understanding the genetic basis of diseases and traits, but most studies have been conducted in isolation, with a focus on either a single or a set of closely related phenotypes. We describe MetABF, a simple Bayesian framework for performing integrative meta-analysis across multiple GWAS using summary statistics. The approach is applicable across a wide range of study designs and can increase the power by 50% compared to standard frequentist tests when only a subset of studies have a true effect. We demonstrate its utility in a meta-analysis of 20 diverse GWAS which were part of the Wellcome Trust Case-Control Consortium 2. The novelty of the approach is its ability to explore, and assess the evidence for, a range of possible true patterns of association across studies in a computationally efficient framework.

## 1 Introduction

In the past decade, a large number of genome-wide association studies (GWAS) have been performed, and they have identified thousands of associations between genotypes and various biological traits (Hindorff et al., n.d.). These associations expand our understanding of both unique and shared molecular mechanisms across different phenotypes (Price, Spencer, & Donnelly, 2015). With so much data available, there is now an increased interest in combining these data in order to learn about regions of the genome that affect multiple traits and, consequently, the shared biological pathways that underlie traits.

While the search for genetic associations shared between traits can be conducted using individual-level genotype data from the participating GWAS (Ellinghaus et al., 2016), in practice, most of the readily available data come in the form of summary statistics. While the exact summary statistics reported vary from study to study, they usually include information about the genetic variant (usually a single nucleotide polymorphism, SNP) and its estimated effect on the phenotype under study. Information about the variant usually includes its genomic location in a given build of the human genome, as well as its alleles, and sometimes the frequencies of the alleles. Information about the association often includes the effect size estimate, its standard error or a 95% confidence interval, and the *p*-value of the association between the variant and the phenotype. The summary statistics information is typically orders of magnitude smaller than the original genotype data files, and reduce the risk of revealing personal information about study participants. For these reasons, a number of methods have been introduced to combine GWAS summary statistics to find genetic loci that influence multiple traits across the genome (Bhattacharjee et al., 2012; Cotsapas et al., 2011; Flutre, Wen, Pritchard, & Stephens, 2013; Majumdar, Haldar, Bhattacharya, & Witte, 2018; Turley et al., 2018; Wen & Stephens, 2014a). These meta-analyses can also find novel associations that were overlooked in individual GWAS.

In this paper we introduce MetABF, an approach to meta-analyse GWAS summary statistics at a single SNP which is simple, efficient, and which can be easily programmed using any standard statistical package—we offer an implementation in R. We have found similar methods to be useful in previous studies (Band et al., 2013; Bellenguez et al., 2012; Gilchrist et al., 2018; Rautanen et al., 2016) and present a unified framework for others to use to explore their data. Unlike most other methods available, ours allows for a direct probabilistic assessment of different models of association across studies, given appropriate prior assumptions. We demonstrate the broad applicability of the approach using simulations of association studies, and by applying it to a diverse set of 20 GWAS that were performed by the Wellcome Trust Case-Control Consortium 2 (WTCCC2, www.wtccc.org). The R implementation is available at https://github.com/trochet/metabf.

## 2 The Method

Traditionally, meta-analysis of GWAS has focused on combining results of multiple studies on the same or similar traits. Figure 1 illustrates the the inference problem. Given a set of effect size estimates 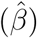, and the uncertainty in these estimates 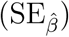 what can we infer about the true effect sizes, *β_i_*, underlying them?

**Figure 1:**
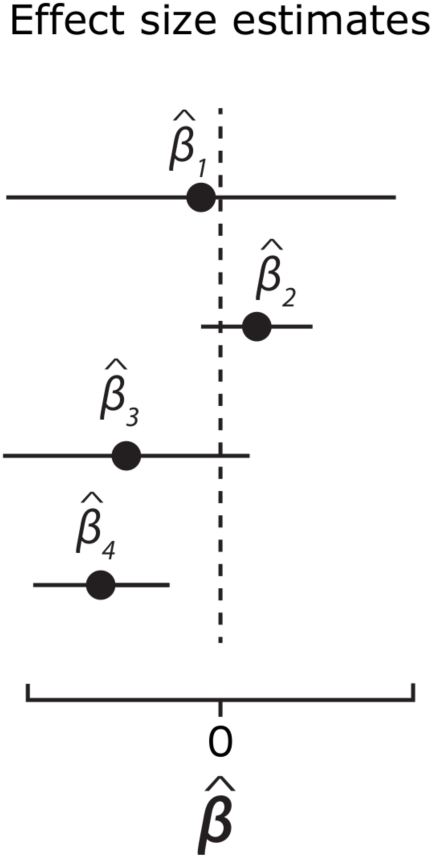
Statement of the problem. Given a set of observed effect sizes estimated from data in *n* = 4 studies, we want to make inferences about the true effect sizes. Our joint estimate of the true effects depends on our assumptions of the similarity between the 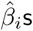 being estimated, for example the similarity between traits being studied. A key idea in our approach is to be able to easily assess the evidence for models with various assumptions of similarity between the studies.

Interpretation of the data depends on assumptions about the heterogeneity in effect between studies. For example, an assessment of the data might be that there is little evidence from studies 1, 2 and 3 because their confidence intervals overlap zero. Even if we were to assume that all studies estimate the same effect, the inconsistency in observed effect sizes, in particular for *β*_2_ and *β*_4_, means that we are unlikely to find strong evidence for a non-zero effect. However, if we knew that the exact trait under examination in studies 1 and 2 was slightly different to those in 3 and 4 then the strong evidence that *β*_4_ *<* 0 might suggest that *β*_3_ *<* 0 as well, without necessarily suggesting the same for *β*_1_ and *β*_2_. Our framework aims to make it easy to quantify the statistical evidence for these kinds of heterogeneous models and to provide the corresponding effect size estimates.

### 2.1 Bayesian approach

Our method uses a Bayesian approach that provides a natural way of capturing the reasoning described above for Figure 1 in a statistical framework. To measure the evidence for association, we calculate Bayes factors, which consider the ratio of the posterior probabilities of two models *M*_1_ (an alternative model) and *M*_0_ (the null model) given some data. By Bayes’ Theorem,

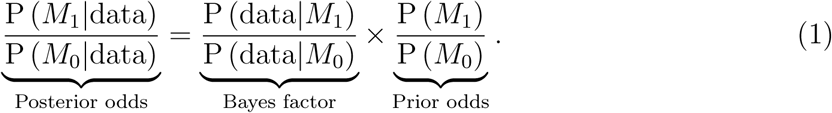

A Bayes factor greater than 1 suggests that the evidence from the data favors *M*_1_ over *M*_0_, while a Bayes factor smaller than 1 suggests the opposite. Note that Bayes factors lie in the range (0, ∞) but when they are presented on a logarithmic scale, negative values are also possible and correspond to the Bayes factors favoring the null model. The interpretation of Bayes factors in the context of GWAS has been discussed earlier (Consortium, 2007; Stephens & Balding, 2009). Because Bayes factors calculate the probability of the data under the null as well as under the alternative model, they naturally account for power to detect effects. In GWAS, this means that Bayes factors can be easier to calibrate across varying study sizes and minor allele frequencies than *p*-values which only consider the tail probabilities under the null model (Wakefield, 2009). Bayes factors require a prior distribution on the model parameters which can be used to describe different alternative models.

In practice, Bayes factors can be computationally expensive to calculate as they commonly involve an integration over the model parameters with respect to the prior distribution. Often, there is no closed form solution of the necessary integrals, necessitating numerical procedures. In the context of GWAS, Jon Wakefield developed an approximate Bayes factor (ABF) that can be calculated directly from GWAS summary statistics (Wakefield, 2007, 2009). Asymptotically, it gives similar results to a Laplace approximation of the integrals in Bayes factor, and the study sizes required for a good approximation are of the order of hundreds of participants (Wakefield, 2009). Here we describe our approach, using approximate Bayes factors (ABFs) in genome-wide association studies (Wakefield, 2007, 2009) which is similar to those applied elsewhere (for example, (Asimit et al., 2015; Pickrell et al., 2016; Wen & Stephens, 2014b)). We compare our MetABF method to two other methods CPBayes (Majumdar et al., 2018) and MTAG (Turley et al., 2018).

### 2.2 Statistical model

For each variant, a GWAS produces a point estimate, 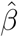, of the effect of a given allele on a trait, typically adjusted for relevant covariates like age, sex, or ethnicity. In quantitative traits, 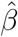 represents the direct effect on the phenotype measurement. In binary traits, 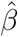 estimates the natural logarithm of the odds ratio (OR) corresponding to each additional copy of the effect allele, that is, exp 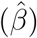 is the odds ratio estimate. Each 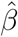 comes with a standard error, 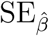, which is estimated along with 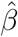 and, in most settings, is largely determined by the sample size and variant frequency. Wakefield’s ABF assumes that given 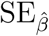, 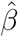 captures the information in the study data about the true effect size, *β* such that

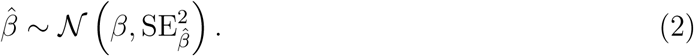

For the large sample sizes found in most GWAS, equation 2 is expected to be a reasonable approximation at all but rare variants (less than 1% frequency). Further discussion of this can be found in section 3.1 of the supplementary material.

Like all Bayesian methods, Wakefield’s ABF includes a prior distribution, which encodes our beliefs about the true effect size *β*. The prior is determined by a single scaling parameter *σ* that is set by the analyst,

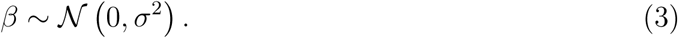

Large *σ* corresponds to a belief that true effect sizes can be large, and analogously for small *σ*. Values commonly used in the literature for disease studies are 0.2 and 0.4 (Marchini & Band, n.d.; Stephens & Balding, 2009). The value *σ* = 0 encodes a belief that the variant has no effect on the trait under investigation, and will be used as the null model, *M*_0_ in our analysis.

For given 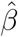, 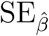, and prior *σ*, let *f* (*x*; *m*, *s*^2^) be the probability density of the Gaussian distribution with mean *m* and variance *s*^2^, evaluated at *x*. Then Wakefield’s ABF is simply

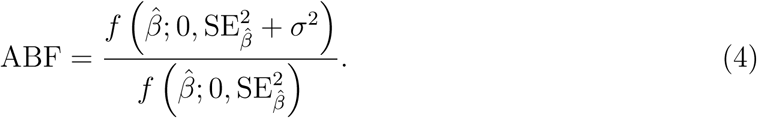

Because this is a ratio of normal densities, it has a closed form expression that can be evaluated very quickly.

We note that the prior distribution on the true effect size, *β*, shown in Equation 3, has most of its probability mass close to zero, which is where the null model has all of its mass. Thus, the Bayes factor in equation 4 has a minimum, non-zero value at 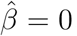, while the maximum value is unbounded. It will rarely provide strong evidence in favor of the null (*M*_0_) as the alternative model includes very small effects close to zero. Other non-local priors (where the prior density of the alternative model drops to zero near zero), have been described (Johnson & Rossell, 2010). The lack of strong evidence in favour of the null reflects the fact that, in the GWAS context, small effect sizes are difficult to rule out. Furthermore, a GWAS typically aims to identify SNPs that show strong evidence (say, ABF *>* 1000) for an association with a trait, rather than those that show strong evidence for no association.

### 2.3 Approximate Bayes factors for meta-analysis

We describe our multivariate extension of the ABF to meta-analysis. Instead of calculating an ABF for an association between a given variant and a given trait measured in one GWAS, we calculate an ABF for an association between a variant and an arbitrary number of traits, *n*, measured in independent or (partially) overlapping studies. Recently, multivariate analysis of GWAS summary statistics has been implemented by the MTAG approach (Turley et al., 2018) and a framework involving Bayes factors to compare models of association has been developed to help determine relevant tissues in eQTL data (Flutre et al., 2013). When extending ABF to the multivariate case, 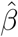, that was a single effect size estimate from a single GWAS, becomes 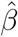, an *n*-vector of effect size estimates from each of the *n* studies included in the meta-analysis.

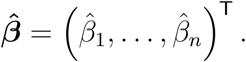

Similarly, 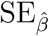 is replaced by the study covariance matrix, 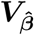, of dimension *n × n*,

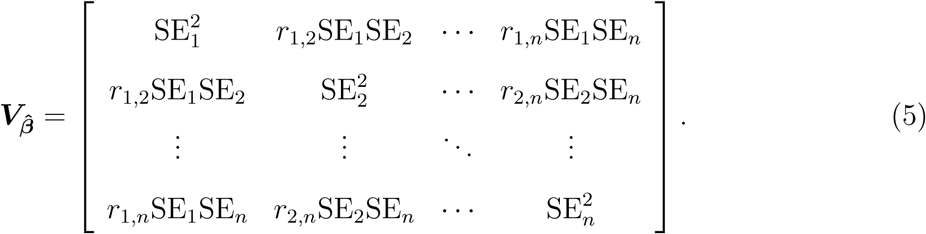

If the studies are all independent—that is, they do not share samples—then the *r_i_*_,*j*_ terms in 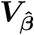 are all 0, and 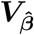 is simply a diagonal matrix of study-wise variances. If studies are not independent, then some or all of the off-diagonal elements of 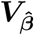 will be non-zero. If the amount of overlap between studies is known, then the values of *r* can be calculated from formulas provided by Zaykin and Kozbur (Zaykin & Kozbur, 2010) and Bhattacharjee *et al.* (Bhattacharjee et al., 2012).

However, having information on exactly which samples were included in which consortia and analyses is increasingly difficult, especially when dealing with summary statistics, necessitating approaches that estimate the covariance between studies directly from the data. The authors of the MTAG approach (Turley et al., 2018) used the intercept of the pairwise LD score regression (B. K. Bulik-Sullivan et al., 2015) to create an analogous matrix. This can be applied to our method as well. As we show later, it is also possible to get similar estimates by simply calculating empirical correlations in a chosen subset of 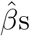 between the two studies.

Next we consider a multivariate extension of *σ*, which we call **Σ**. As in the univariate case, each study *i ∈ {*1, …, *n}* has a prior parameter *σ_i_* determining the expected effect sizes. However, combining multiple studies introduces the possibility of correlated true effects between the studies, which we capture in the terms *ρ_i_*_,*j*_: *i*, *j ∈ {*1, …, *n}*.

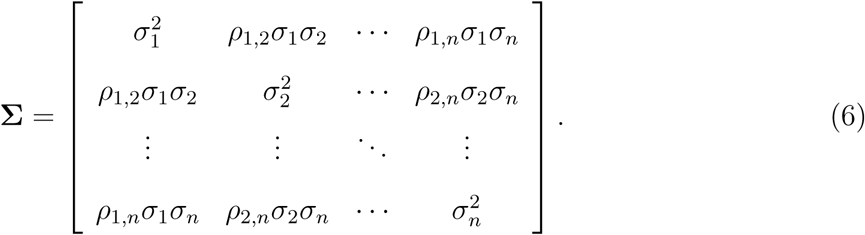

This matrix has a similar role as the **Ω** matrix in the MTAG approach (Turley et al., 2018). Thus, by writing *f* (***x***; ***m***, ***V***) for the density of a multivariate normal distribution with mean vector ***m*** and covariance matrix ***V***, evaluated at ***x***, Equation 4 becomes

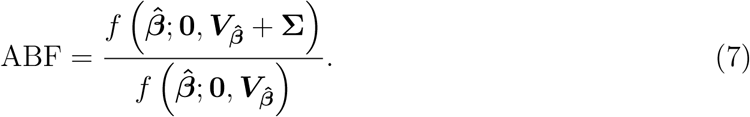

### 2.4 Choosing prior parameter values

Unlike 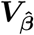, which is in theory can be defined by the data, **Σ** reflects the prior beliefs of the researcher about the similarity of the effects between studies. One of the most common ways to meta-analyze a set of GWAS summary statistics is the inverse-variance weighted fixed-effects model. This model assumes that the underlying true effect of an associated variant is the same in all cohorts, and the differences in the estimated effect sizes arise due to statistical noise. Larger studies produce more accurate estimates of the underlying true effect producing estimators with smaller variances. Thus, weighting by the inverse of the variance gives a larger study a greater contribution to the meta-analyzed estimate than a smaller one. In our framework, this fixed-effects model is encoded by setting all the *ρ_i_*_,*j*_ terms of **Σ** equal to 1.

Another common meta-analysis method is the random-effects model. This assumes heterogeneity in true effect sizes across studies. There are many ways of implementing this assumption, many of which have inherently reduced power to detect effects compared to the fixed effects approach (Han & Eskin, 2011). Our approach allows for a number of different priors that allow for heterogeneity in effects found in different studies. They are summarized, along with the fixed-effects prior, in Table 1.

**Table 1:** Different models of association across studies and their relation to parameters of the prior model. Here σ and *ρ* are as defined in text and assumed to be the same across studies.

|  | Variance of<br>effect size | Correlation<br>in effect<br>sizes | Notes |
| --- | --- | --- | --- |
| Null model | $\sigma^2 = 0$ | - | Assumes variant has no effect on the phenotype. |
| Fixed effects | $\sigma^2 > 0$ | $\rho = 1$ | Equivalent to first applying an inverse-variance weighted meta-analysis to estimates $\hat{\beta}_i$ and $SE_{\hat{\beta}_i}$, and then using these estimates as a single study in equation (4). |
| Independent effects | $\sigma^2 > 0$ | $\rho = 0$ | Equivalent to multiplying the single study ABFs across studies, assuming the studies are independent. Similar to Fisher’s method of combining $p$-values. |
| Correlated effects | $\sigma^2 > 0$ | $0 < \rho < 1$ | Similar to a related-effects approach where the effects are assumed to be correlated after being drawn from a common distribution (see text). |
| Subset model | $\sigma_i^2 > 0$ for $i$<br>in a subset<br>$I \subseteq \{1, \dots, n\}$ | $0 \leq \rho_{ij} \leq 1$<br>for $i, j \in I$ | Models heterogeneous patterns of effects, where only a subset of the underlying parameters are nonzero. May be most appropriate to model as fixed, independent, or correlated within the set of nonzero effects, depending on the setting. |

The prior models described above allow us to quantify the evidence of association through an ABF in which the alternative model can take a range of forms. The simplest models assume that all studies draw their effects from the same marginal distribution and are equally correlated with one another, such that the prior can be specified with just a single *σ* and a single *ρ* parameter. When *ρ <* 1, the prior allows some studies to have large effects while others can still have effects very close to zero since the prior on *β* is always centered on zero. In this way, we expect an ABF which assumes all studies have an effect with 0 *< ρ <* 1 to show evidence for the alternative model, that is, ABF*>* 1, even when only a subset of the studies have truly nonzero effects.

### 2.5 Subset exhaustive model averaging

In the analysis of a diverse set of phenotypes, it is natural to consider the possibility that only a subset of studies have non-zero effects 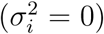 while the rest are null 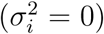. For *n* studies and fixed values of *ρ* and *σ*, there are 2^*n*^ possible subset models, including the null model of no association. A natural Bayesian approach for detecting associations under these assumptions is to apply model averaging over the subset models to obtain a single summary of the evidence for models of association against the null model of no association. By storing the ABF for each of the models, the analysis can also generate the full posterior distribution over the models. These analyses require specifying a prior probability on each model. When all studies are considered equally likely to have an effect, then possible prior distributions include:

**Uniform prior:** This assumes that every model of association is equally likely, and thus has the prior probability *p* of

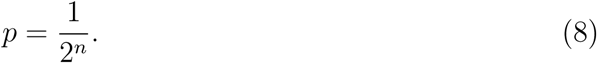

**Combinatorial prior:** When calculating the total prior weight on the models with *m* associated traits out of all possible *n* traits under the uniform prior, we see that the models with 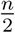 traits (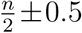 if *n* is odd) have the largest prior weight since the number of these models is the largest. The combinatorial prior forces the prior weights on the number of associations to be the same. That is, for a model with a total of *m* associations, the prior probability *p* on the model is

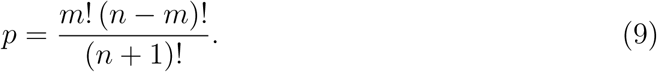

**Binomial prior:** Both the uniform prior and the combinatorial prior assume that a model with no associations is equally likely as a model where all *n* traits are associated. In reality, we might assume that each of the studies has some probability *q_i_*: *i ∈ {*1, …, *n}* of having a true association with a variant. Alternatively, these *q_i_* can be thought of as the proportion of SNPs with which the *i*th study is expected to be associated. This is similar to *π*, the prior on the expected proportion of true effects that Stephens and Balding (Stephens & Balding, 2009) discuss for the calculation of the posterior probability of association. For a given model with subset *K ⊆ {*1, …, *n}* of studies showing associations with the variant and subset *L* = {1, …, *n}\ K* showing no association, the prior probability of the model is

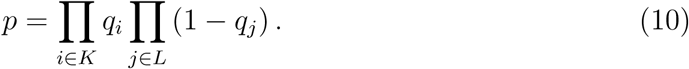

The uniform prior can be recreated in this prior by setting all *q_i_* = 0.5.

In practice, we do not recommend using the combinatorial prior, as it tends to result in the highest ABF being associated with either the null model or the model with all traits associated. This becomes more pronounced as the number of studies in the meta-analysis, *n*, increases. This is because for all values of *n >* 1, there is exactly one null model and one model where all *n* traits are associated, while the number of models with one to *n −* 1 associations increases with *n*, so the prior weight on any one of these models decreases with *n*, which also decreases their relative weights compared to the null and *n* associated models. Furthermore, in real world applications, it is unlikely that one would have an *a priori* belief in the number of true underlying associations while being completely agnostic to which specific traits are associated.

By making use of the generality of the prior covariance matrix **Σ**, a range of other models can be defined based on particular assumptions on the relationship between traits. Where it might be appropriate to assume that the marginal effect at a variant is representative of the genome average, the estimates of the genetic correlation between two studies might be appropriate as prior information, and it has been shown that this can increase power in some scenarios (Turley et al., 2018).

In practice, for analyses of large numbers of studies, we advocate selecting a relatively small set of prior matrices which best represent possible models as a way to overcome the exponentially growing model space and the fact that prior probability on any one subset model becomes increasingly small with increasing number of studies. The rationale is similar to the approach taken in the analysis of eQTL data (Flutre et al., 2013). We can average across all of the ABFs calculated under the various priors to get a single quantity that accounts for our uncertainty about the exact prior model.

It is also of interest to assess models in which effects are assumed to be negatively correlated between some pairs of studies. A challenge is that the number of possible configurations of positive and negative association grows even more quickly than the subset models. Taking absolute values of the effect sizes changes the sampling distribution to the folded-Gaussian distributions which are difficult to work with. If negative correlations are of interest, or should be entertained as possible, then assuming independent effects (*ρ* = 0) is a practical option, although with the cost of losing the dependency information between the effect sizes.

#### 2.5.1 Shotgun stochastic search

If the number of studies in the meta-analysis is so large that an exhaustive search over all subset models is not feasible for a given **Σ**, a shotgun stochastic search (SSS) (Hans, Dobra, & West, 2007) can be performed to determine the most likely models of association. This iterative method calculates the ABFs for the set of models in the “neighborhood” of the current index model. This neighborhood contains models that (i) add an association to the index model, (ii) remove an association from the index model, or (iii) replace exactly one nonzero study from the index model with one that is zero in the index model. The algorithm then chooses the next index model based on a probability distribution generated from normalizing the ABFs of the models in the neighborhood of the index model. This type of search has been used to fine-map genetic loci (Benner et al., 2016) and is well-suited to our ABF analysis, as it quickly finds the models with high ABFs. In every scenario we have investigated, in both real and simulated data, the distribution of ABFs has been unimodal, meaning that the search is unlikely to miss the global maximum of the distribution due to being caught in a local one.

### 2.6 Posterior distribution of effect sizes

Within the Bayesian framework described above, it is natural to compute the posterior on the true effect sizes under different assumptions about the presence and size of an effect within a study, and the correlation in effects across studies. Often when describing the interpretation of a forest plot, like that illustrated in Figure 1, the analyst informally imposes these beliefs in drawing conclusions. It is therefore of interest to assess formally the impact of these beliefs on our inference about the true effects.

Using standard results for the posterior distribution of the mean given a multivariate normal likelihood and a normal prior:

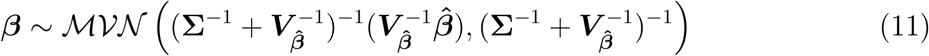

A posterior forest plot can then be drawn with point estimates based on the posterior mean, and with the marginal variance on the posterior effect size taken from the diagonal elements of the posterior variance-covariance matrix. The impact of the prior matrices described above on the posterior forest plot typically includes: shrinkage of effects towards zero, when they are believed to be non-zero; increased certainty in the effect size estimates of two studies with similar effects when they are assumed to be correlated, with the certainty at its maximum when the studies are independent; and increased uncertainty in true effects which are assumed to be correlated when the observed effects differ significantly. We note that it would be possible model average the posterior distribution over models by numeric evaluation. In this case, the marginal posterior distribution of effect size within a study is no longer necessarily unimodal.

## 3 Results

### 3.1 Simulations and statistical properties

To understand the statistical properties of our method compared to standard approaches to meta-analysis, we performed simulations of 5 independent genome-wide association studies at a single variant. We found that, assuming sample sizes are moderate (*>* 1000 individuals) and allele frequencies not too rare (*>* 0.01), it was possible to simulate effect sizes and standard errors directly from the the model described above, where true effects can either be fixed at a given value, or sampled from a prior distribution (see Supplementary Material 6.3). The ability to efficiently simulate directly from the approximate model is useful for assessing frequentist properties of the Bayesian approach.

As an illustrative example Figure 2 shows the estimated power of a standard inverse-variance weighted fixed effects meta-analysis approach and Fisher’s method of combing *p*-values, as well the power of a MetABF which assumes that all studies have an effect. Results are shown for MetABFs that assume effects are uncorrelated (*ρ* = 0), or highly correlated (*ρ* = 0.96). The MetABF with highly correlated effects performs well when there is an effect in all 5 studies, although not quite as well as the fixed effects approach when the fixed effects assumption is correct. However it has substantially more power when only a subset of studies—for example 2 out of 5—have an effect in which case the highly correlated MetABF has 50% more power than the fixed effects approach.

**Figure 2:**
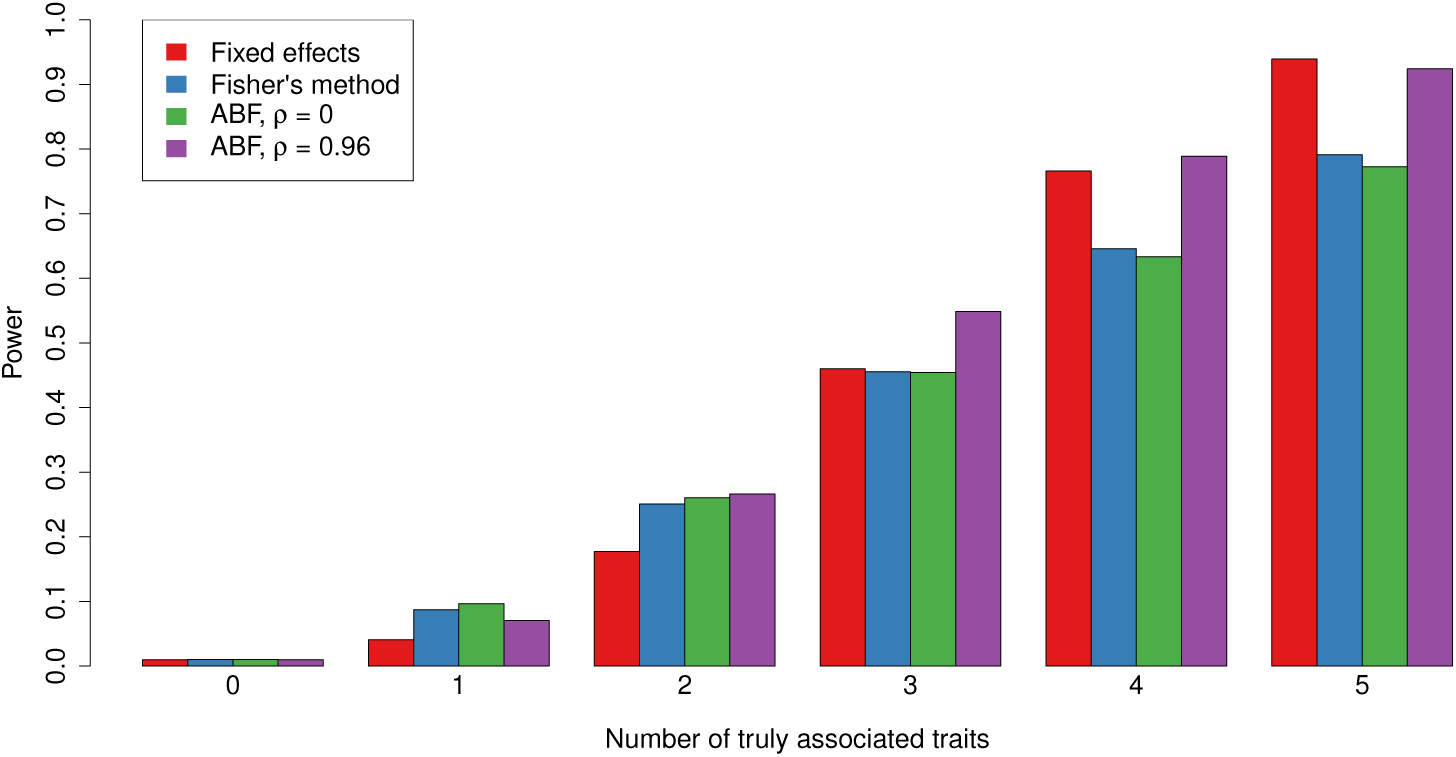
Plot of power for two frequentist test and the ABF with two different prior parameter settings for the assumed correlation between studies (*ρ*). Simulation were performed with a true effect (Odds ratio = 1.1) present in a subset (y-axis) of 5 studies with 1500 cases and 1500 controls. Power was calculated using a significance threshold *α* = 0.01. ABFs were calculated with *σ* = 0.2.

We used simulations under the null to estimate *p*-values for the ABF using priors which assumed effects were present in all 5 studies (as in Figure 2), and the MetABF model that explicitly averages over all possible subsets of true effects. We note that when the ABF is based on a single variance-covariance matrix for the prior, i.e. it is not model averaged, then it is possible to derive the distribution of ABF under the null, given a frequentist test statistic, see Supplementary Material 6.5. We performed simulations under a range of different subset effects with different priors on effect sizes and correlations and compare statistical power to two frequentist methods described above as well as a random effects approach with a closed-form calculation (DerSimonian & Laird, 1986).

Briefly, we found that the MetABF approach was similar in power to the best frequentist tests, was more powerful when a subset of the traits in the meta-analysis were truly associated. Of the three alternative approaches we tested, Fisher’s method came closest in terms of power with the Bayes factor approaches when there are three or more studies with associations to find. We also found that while MetABF sometimes loses power when the prior correlation on the true effects is different from the true underlying correlation, the loss is not large — the results of the MetABF analyses using different values of *ρ* (but the same prior *σ*) tended to differ by less than 0.005. This suggests that the MetABF approach is fairly robust to the choice of prior *ρ*.

### 3.2 Comparison to other methods

We compared our method to both CPBayes (Majumdar et al., 2018) and MTAG (Turley et al., 2018) using simulated data. Section 7 in the supplement provides a more detailed discussion of our findings. Briefly, at any given false positive rate (proportion of spurious associations in the posterior model), CPBayes had the highest true positive (proportion of true associations in the posterior model) rate of the three methods, while MetABF and MTAG were very similar in performance. However, neither MTAG nor CPBayes allow the user to explore the posterior probability space the way MetABF does. Additionally, the gain in accuracy in CPBayes comes at a considerable time cost. It took roughly 8 minutes on average to analyze a dataset of 60,000 SNPs with MetABF, while CPBayes took an average of 2.3 days when studies were assumed to have no cryptic relatedness between them, and an average of 6.6 days when they were not. Both CPBayes and MetABF are implemented as R packages. MTAG, which is implementedin Python, was by far the fastest method, taking an average of 15 seconds on the full data; however, this does not include the time taken to create the LD score reference panels.

### 3.3 Application to WTCCC2 data

To investigate the applicability of the approach across a large number of heterogeneous GWAS, we applied it to summary statistic data from the following Wellcome Trust Case Control Consortium 2 (WTCCC2) studies. These included

**Autoimmune diseases:** 1) ankylosing spondylitis (AS); 2) multiple sclerosis, UK cohort (MS UK); 3) multiple sclerosis, non-UK European cohort (MS nonUK); 4) psoriasis (PS); 5) ulcerative colitis (UC).

**Infectious diseases:** 6) bacteremia, all types (BS overall); 7) pneumococcal bacteremia (BS pneumococcus); 8) visceral leishmaniasis, Indian cohort (VL India); 9) visceral leishmaniasis, Brazilian cohort (VL Brazil).

**Stroke cohorts:** 10) ischemic stroke, large vessel subtype (IS TOAST 1); 11) ischemic stroke, small vessel subtype (IS TOAST 2); 12) ischemic stroke, cardioembolic subtype (IS TOAST 3).

**Reading and mathematics cohort:** 13) reading scores (RM reading); 14) mathematics scores (RM maths).

**Psychiatric traits:** 15) schizophrenia (SP); 16) psychosis endophenotypes (PE).

**Studies of unique traits:** 17) Parkinson’s disease (PD); 18) Barrett’s esophagus (BO); 19) metformin response (PR); 20) glaucoma (GL).

Studies were mainly conducted in European cohorts, but two studies (BS and VL) used non-Europeans samples. Three different genotyping arrays were used, and four studies were imputed to increase the number of SNPs in the dataset (see Supplementary Tables 7 and 8 for details). As a result, summary statistic data were typically available for only a subset of studies at any given SNP. Our analysis included all SNPs for which there were summary statistics in at least two of the studies. Before performing our analysis we harmonised the data using a pipeline to align the SNPs in each study to the forward strand, thus ensuring that the “A” and “B” alleles of each SNP were the same across all studies, and corresponded to the reference/alternative alleles in the 1000 Genomes database. Effect sizes were estimated for the alternative allele. Details about the processing and availability of the data can be found in Section 6.7.1 of the Supplementary Material.

To search for signals of association we calculated a model averaged ABF at each SNP across the genome. To capture the range of possible patterns of association across studies, we selected 12 models with different assumptions about the correlation structure of effect sizes across studies. These are described in section 3.4 below. We found that subset models that assumed an effect only within one study (and explicitly no effect in other studies) often had low probability, presumably because conditional on one study showing a true effect, the chance that no other study shows any effect becomes increasingly small (particularly for related phenotypes), as the number of studies increases. We calculated the ABF for each of these models with different values of *σ* (0.1,0.2 and 0.4). We then took the mean across all resulting ABFs, assuming them to be equally likely a priori, to obtain a single summary of the association evidence across all studies.

### 3.4 Prior matrices

Briefly, the 12 priors used in our analyses assume non-zero effects across all studies. They differ in the presumed correlation structure, ranging from completely uncorrelated across all studies (model 1), through to prior matrices where correlated effects only occur within a subset of studies (for example IS, MS, BS and RM, models 5-7) or within study classes (for example autoimmune disease; AS, PS, MS and UC: models 8-12). Figure 3 illustrates each of the prior models used. The traits are listed in the order described at the start of Section 3.3, grouped by disease/trait class.

**Figure 3:**
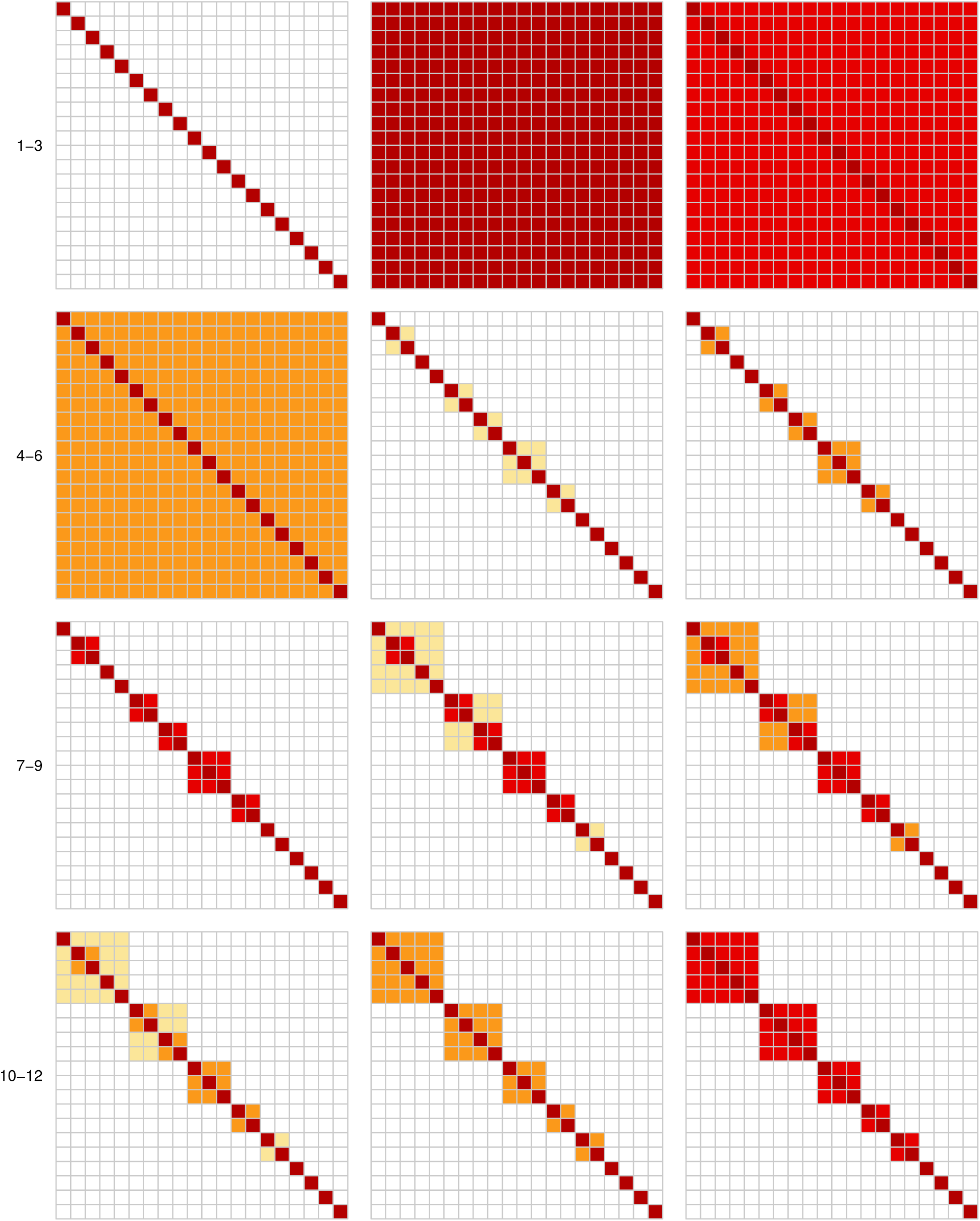
Visualizations of the prior correlation matrices of WTCCC2 analyses. Dark red squares correspond to *ρ* = 1, red squares correspond to *ρ* = 0.96, yellow-orange squares correspond to *ρ* = 0.5, light yellow squares correspond to *ρ* = 0.1, and white squares correspond to *ρ* = 0. The traits are listed in the order at the start of Section 3.3 and grouped by disease/trait class.

### 3.5 Study matrix

Among the 20 WTCCC2 cohorts, nine of them used samples from a shared pool of controls. Additionally, one of the bacteremia (BS) cohorts was a subset of the other, while the two reading and mathematics cohorts (RM) comprised of two different phenotypes measured on the same individuals. This introduced correlation in the effect size estimates due to shared statistical noise, and therefore non-zero covariance terms between some of the studies in the 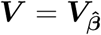 matrix (see Equation 5), which we needed to estimate (Table 2).

**Table 2:** Comparison of the 39 non-zero correlations calculated under the null between each pair of studies using different methods: using the set of SNPs shared between the two phenotypes with genome-wide significant SNPs and those in the MHC region removed (“Signals removed”), the set of SNPs shared between the two phenotypes, thinned so that all SNPs are at least 0.25 centiMorgans apart (“Thinned”), the set of all SNPs shared between the two phenotypes (“All”), and the intercept of the genetic covariance calculated by LD Score regression B. Bulik-Sullivan et al. (2015); B. K. Bulik-Sullivan et al. (2015) (“LDSC”).

| Trait 1 | Trait 2 | Signals |  |  |  |
| --- | --- | --- | --- | --- | --- |
|  |  | removed | Thinned | All | LDSC |
| AS | BO | 0.192 | 0.162 | 0.177 | 0.196 |
| AS | IS TOAST 1 | 0.097 | 0.0946 | 0.0932 | 0.0939 |
| AS | IS TOAST 2 | 0.0995 | 0.0892 | 0.0868 | 0.0998 |
| AS | IS TOAST 3 | 0.0885 | 0.0817 | 0.0804 | 0.096 |
| AS | MS UK | 0.184 | 0.143 | 0.147 | 0.199 |
| AS | PD | 0.198 | 0.185 | 0.169 | 0.206 |
| AS | PS | 0.198 | 0.147 | 0.152 | 0.206 |
| AS | UC | 0.213 | 0.203 | 0.209 | 0.225 |
| BO | IS TOAST 1 | 0.122 | 0.134 | 0.122 | 0.127 |
| BO | IS TOAST 2 | 0.133 | 0.162 | 0.132 | 0.135 |
| BO | IS TOAST 3 | 0.113 | 0.121 | 0.113 | 0.126 |
| BO | MS UK | 0.229 | 0.199 | 0.226 | 0.241 |
| BO | PD | 0.237 | 0.227 | 0.237 | 0.236 |
| BO | PS | 0.284 | 0.277 | 0.273 | 0.294 |
| BO | UC | 0.244 | 0.251 | 0.244 | 0.268 |
| BS overall | BS pneumo. | 0.619 | 0.623 | 0.619 | NA |
| IS TOAST 1 | IS TOAST 2 | 0.121 | 0.133 | 0.120 | 0.127 |
| IS TOAST 1 | IS TOAST 3 | 0.169 | 0.175 | 0.168 | 0.164 |
| IS TOAST 1 | MS UK | 0.115 | 0.098 | 0.111 | 0.118 |
| IS TOAST 1 | PD | 0.111 | 0.118 | 0.111 | 0.110 |

|  |  | **Signals** |  |  |  |
| --- | --- | --- | --- | --- | --- |
| **Trait 1** | **Trait 2** | **removed** | **Thinned** | **All** | **LDSC** |
| IS TOAST 1 | PS | 0.129 | 0.119 | 0.124 | 0.127 |
| IS TOAST 1 | UC | 0.114 | 0.112 | 0.115 | NA |
| IS TOAST 2 | IS TOAST 3 | 0.117 | 0.137 | 0.118 | 0.114 |
| IS TOAST 2 | MS UK | 0.124 | 0.127 | 0.121 | 0.127 |
| IS TOAST 2 | PD | 0.128 | 0.117 | 0.127 | 0.128 |
| IS TOAST 2 | PS | 0.134 | 0.136 | 0.127 | 0.139 |
| IS TOAST 2 | UC | 0.124 | 0.161 | 0.123 | 0.128 |
| IS TOAST 3 | MS UK | 0.112 | 0.0866 | 0.11 | 0.116 |
| IS TOAST 3 | PD | 0.112 | 0.102 | 0.112 | 0.104 |
| IS TOAST 3 | PS | 0.113 | 0.108 | 0.107 | 0.122 |
| IS TOAST 3 | UC | 0.109 | 0.131 | 0.109 | NA |
| MS UK | MS nonUK | 0.0529 | 0.132 | 0.0927 | 0.0191 |
| MS UK | PD | 0.243 | 0.245 | 0.242 | 0.252 |
| MS UK | PS | 0.243 | 0.218 | 0.222 | 0.254 |
| MS UK | UC | 0.233 | 0.224 | 0.231 | 0.244 |
| PD | PS | 0.249 | 0.269 | 0.237 | 0.253 |
| PD | UC | 0.24 | 0.251 | 0.239 | 0.246 |
| PS | UC | 0.257 | 0.247 | 0.240 | 0.271 |
| RM reading | RM maths | 0.535 | 0.535 | 0.535 | 0.543 |

When estimating the correlation directly from the genome-wide 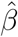-statistics, we used several approaches: taking the intercept from LD score regression (LDSC) (B. Bulik-Sullivan et al., 2015; B. K. Bulik-Sullivan et al., 2015; Turley et al., 2018), which provides an estimate of trait covariance due to undetected shared controls; using all the available data and empirical correlation of 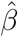-estimates over three subsets of the SNPs (all SNPs; thinning out the SNPs by the recombination distance; and removing SNPs that had significant associations with *p <* 5 × ^-10^ as well as those in the MHC region). Broadly, the estimated correlations were similar across approaches, however the removal of significant associations and the MHC region tended to increase the estimated correlation, as did the LDSC analysis. These observations suggest that strong association signals create additional variance in the effect size estimates that can reduce the correlation if not accounted for.

For the analysis that follows, we assumed ***V***_*i,j,p*_ = *r*_*i,j*_*SE*_*i,p*_*SE*_*j,p*_, where *i* and *j* indexed the study pairs and *p* indexed the SNP. Because LD score regression was not able to run on all of our data, due to small (*<*100,000) overlaps between the markers in software’s reference panels and the markers in some of our cohorts, the pairwise correlations *r_i_*_,*j*_ were based on the genome-wide correlations in 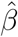 estimated after the removal of the MHC region and other significant associations, and *SE_j_*_,*p*_ were taken directly from the original association analysis.

Due to simplicity of the multivariate normal approximation, it is quick to simulate effect size estimates and standard errors under the null for all 20 traits in our analyses. Using these simulated data, we can calculate ABFs under a range of models, across all traits, and simultaneously accounting for the non-independence of the studies. It is then possible to check, via a quantile-quantile comparison, whether the observed ABFs are distributed as expected under the null at the majority of SNPs, which would provide confidence that positive ABFs represent evidence for genuine departures from the null model, as opposed to systematically mis-calibrated test statistic. Supplementary Figure 13 shows the result of the simulations across the genome for each of the 12 prior models. The resulting quantiles are closely matched at the majority of SNPs suggesting that the null model fits the data at the majority of SNPs. As expected there is a deviation in the tail, where the observed ABFs are bigger than expected, suggesting the alternative model is a better explanation for the data. We caution that in the analysis of genome-wide association data, these simulations do not account for the correlation between SNPs due to linkage disequilibrium.

### 3.6 Genome-wide analysis

The results of the genome-wide analysis in Figure 4 show the mean MetABF at each SNP across all 36 combinations of 12 prior matrices and three values of *σ* (*σ* = {0.1, 0.2, 0.4}), using the model that assumes an association in every trait. Additionally, we curated two lists: one of SNPs reported in each WTCCC2 publication as being implicated by previous studies, and the other of novel loci identified as genome-wide significant (often after replication) by each publication. These are given in the Supplementary Material (Tables 9 and 10). Of the 35 previously identified associations, 18 had a model averaged MetABF *>* 10^4^ in our analysis.

**Figure 4:**
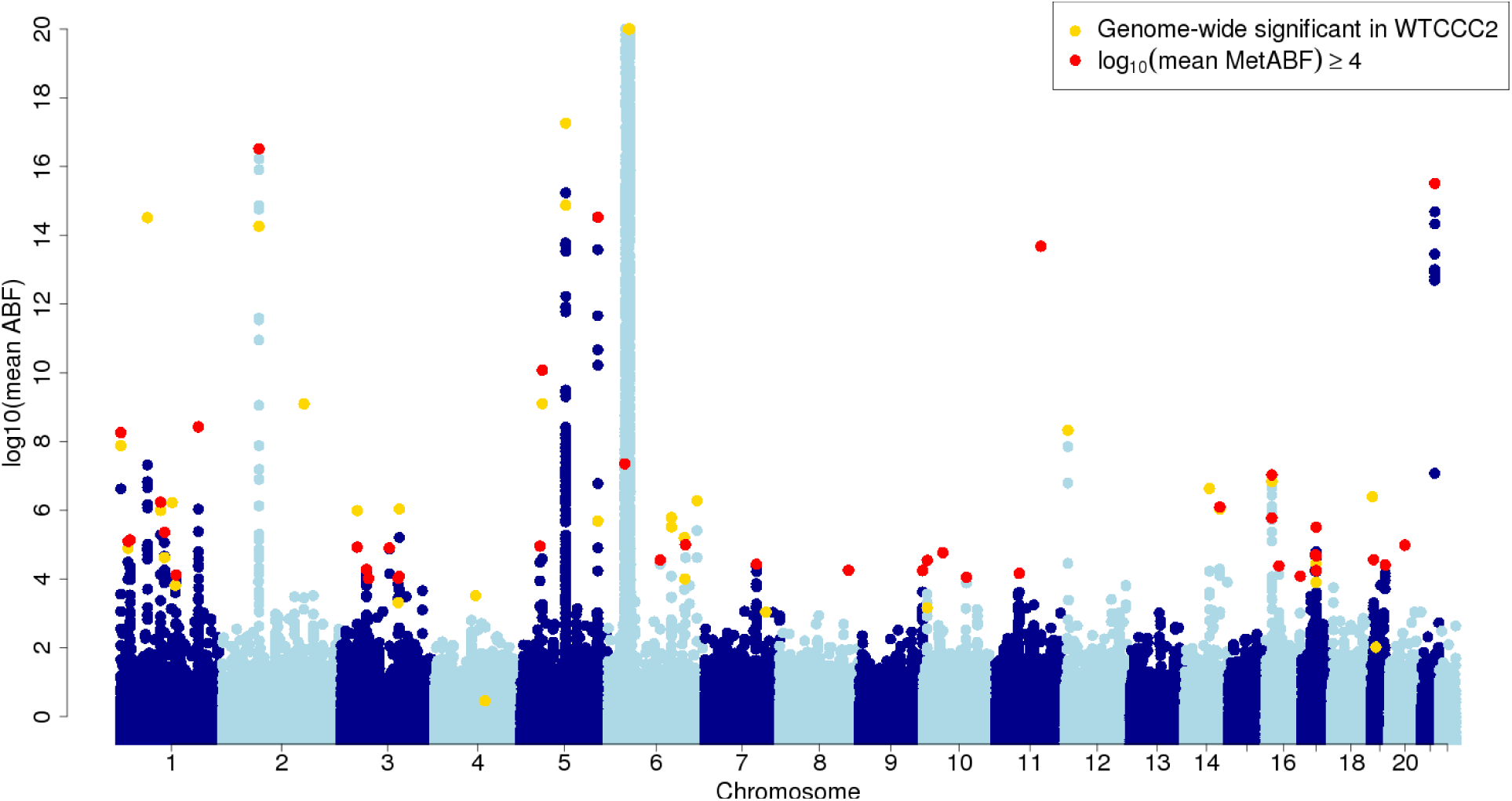
MetABF analysis of genome-wide association analysis across 20 WTCCC2 studies. Adjacent chromosomes are coloured with different shades of blue with SNPs of interest colored as denoted in the legend. The y-axis has a threshold at 20. We highlight the markers with the highest mean ABF in our analysis for each non-MHC region with markers whose mean MetABFs were greater than 10^4^. Yellow dots show markers that had been established as loci associated with one or more of the WTCCC2 traits. Red dots highlight the SNPs with the highest mean MetABF (for a given region) that show an association with at least one trait in the meta-analysis, where the mean MetABF ≥ 10^4^, and which are not genome-wide significant in any of the original WTCCC2 analyses.

For regions showing a strong evidence of association (model-averaged MetABF *>* 10^4^) we further explored the patterns of association, examining all possible subset models across all 12 prior correlation matrices and three values of prior effect size parameter *σ*. For markers with information for at least 19 studies, we used a shotgun stochastic search (Hans et al., 2007) to avoid making all 2^19^ or 2^20^ calculations. Because of the large number of possible models, and because we assumed each to be equally likely a priori (Uniform prior), this approach imposes a strong prior belief in the models with an intermediate number of non-zero effects. We show the marginal probability of the study being included (averaged across all possible subset models) and the most likely model in Supplementary Figure 14. Outside of the MHC region (the strong peak of association on chromosome 6), several loci stand out as potentially having effects across a range of traits. To illustrate the impact of the ABF analysis on the interpretation of the pattern of association we highlight three SNPs. The forest plots of the original association summary statistics and the posterior distribution of effects under the most likely model are shown in Figure 5. We discuss these illustrative examples below.

**Figure 5:**
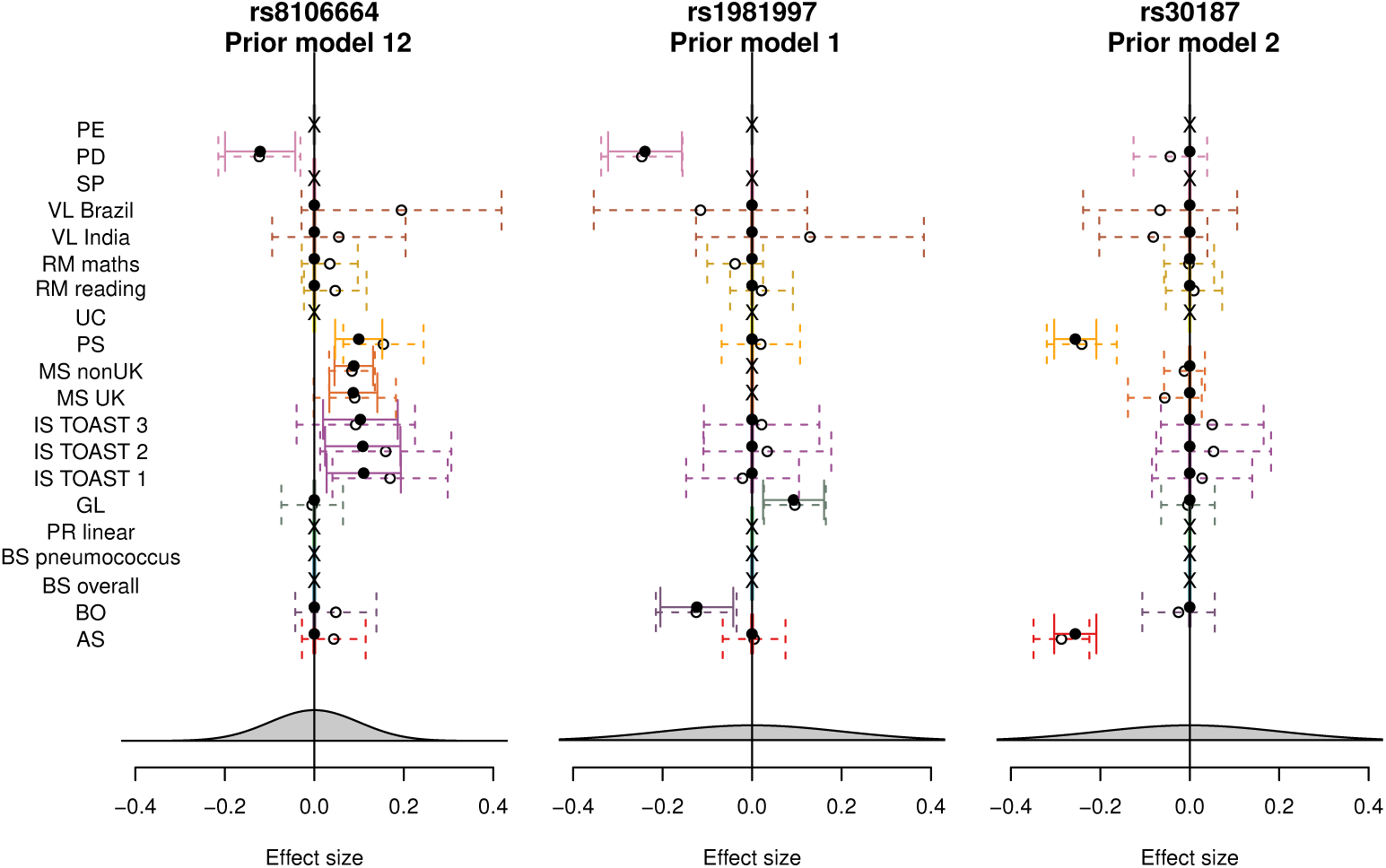
Bayesian forest plots of three SNPs highlighted in the genome-wide scan under the prior assumption for the most likely posterior model. The dotted lines and open circles show the original effect size and 95% confidence intervals in the WTCCC2 studies. The filled circles and the solid lines show the mean and 95% credible regions from the posterior distribution under the top model (crosses indicate the SNP was missing). Posterior effects at zero reflect that the top model assumed no effect in the study. Grey densities at the bottom show the prior distribution on the effect size under the top model.

- *SLC44A2* locus - rs8106664. The analysis highlights a SNP near the gene *SLC44A2* which shows strong association with MS, as well as associations with both psoriasis and ischemic stroke. The SNP was not identified in the 2011 multiple sclerosis analysis (Sawcer et al., 2011), but is in strong LD (*r*^2^ = 0.9347 in 1000 Genomes EUR populations) with a missense variant that was subsequently confirmed (Beecham et al., 2013). Its prominence in this analysis is driven by the signals in other auto-immune diseases which boost the signal in MS and increases the confidence in the effect size. The most likely model also includes an effect in Parkinson’s disease, which under the prior correlation matrix (prior model 12, which assumes high correlations among traits from the same class, but no correlations otherwise) is independent of the other studies, allowing it to have an effect in the opposite direction. *SLC44A2* is upstream of *LDLR*, a well known lipid locus, and downstream of *TYK2*, which has a complex association with multiple auto-immune diseases. One possibility is that this SNP is linked to multiple signals of association and therefore is not necessarily causal one for every, or even any, of the traits showing association (see Supplementary Figure 15). In any case, this analysis highlights a genetic variant that is a marker for susceptibility to multiple traits.
- Chromosome 17 inversion - rs1981997. There is a large polymorphic inversion on chromosome 17 (Stefansson et al., 2005; Tobin et al., 2008) which contains several important genes, including the gene *MAPT* which shows a strong signal of association with Parkinson’s disease (PD) (Skipper et al., 2004; Steinberg et al., 2012; Tobin et al., 2008). A number of SNPs which tag the inversion show strong evidence of association in our analysis. The size of the effect in PD leads to the top model having an effect size prior with a variance of 0.2^2^, which increases the plausibility of large effects in other studies. Interestingly, the top models include associations in glaucoma in the opposite direction to the PD risk, and Barrett’s oesophagus (BO) showing effects in the same direction with PD.
- *ERAP1* mutation - rs30187. The strongest signal of association outside of the MHC was at rs30187, which was previously reported as being significantly associated with both PS and AS, and these effects drive the evidence for association also in our analysis. The variant is a missense change (Lys528Arg) with the effect allele associated with impaired peptide trimming, which is more strongly protective in the presence of a pre-disposing HLA allele (Evans et al., 2011). The top model identifies this effect as being specific to AS and PS. This model also assumes effect sizes are strongly correlated between all diseases that have an effect, with a prior variance of 0.2^2^. These assumptions couple the effect size estimates, marginally reducing the posterior estimates of the study-wise standard errors for AS and PS, and increase the effect in PS to become more consistent with AS.

The above examples highlight how the subset models can explicitly quantify the extent to which that the data are consistent with no effect at all in a subset of studies, and assess assumptions about the size and correlation in true effects across the remaining studies. The analysis explores the posterior probability on 2^20^ × 12 × 3 = 37,748,736 models, and therefore has substantial flexibility to reveal patterns from the data. An advantage of this analysis is that any two models can be compared directly against each other to explore how well the data support one model over the other.

## 4 Discussion

Here we have introduced MetABF, a method for searching for cross-trait associations using GWAS summary statistics. The Bayesian approach allows the expected relationships between studies or traits to be encoded in the analysis. When effects are assumed to be correlated between studies, a strong effect in one study in the meta-analysis automatically adjusts the threshold of evidence required to discover extra associations among related studies. Because the calculations are fast, we can compare different models of association under different priors directly to determine which fits the data best. This allows us to make probabilistic statements about which studies showed true effects at a given marker.

This type of approach is attractive in its ability to combine a large number of heterogeneous GWAS, so to demonstrate how it can be applied, we jointly analyzed 20 different traits across the WTCCC2 data. Our results highlighted loci that showed multiple associations across traits studied which did not achieve genome-wide significance in the individual GWAS. Two different genotyping protocols were used across all the WTCCC2 studies, and some studies contained imputed markers, meaning that there were data for all traits at fewer than 5% of the markers in our meta-analysis. In general, our method would tend to favour a SNP for which there is information on more studies over a SNP for which there is less data, but which shows a slightly stronger effect in one of the studies.

Cross-trait analyses, including ours have limitations. Specifically, markers with apparent cross-study associations may be tagging multiple distinct causal loci. This is a problem inherent in GWAS in humans, where the typed marker is rarely the causal variant, but in linkage disequilibrium with it (Visscher, Brown, McCarthy, & Yang, 2012). Implicitly this up-weights SNPs that tag multiple causal SNPs and complicates direct interpretation of the patterns of associations across studies. Even when the same typed marker is significant for multiple GWAS, the underlying causal markers may be different, and this may be exacerbated by differences in the patterns of linkage disequilibrium between population under examination (Giambartolomei et al., 2014). Colocalization methods (Giambartolomei et al., 2014; Hormozdiari et al., 2016; Wen, Pique-Regi, & Luca, 2017) may be employed to determine if multiple causal variants are likely for regions that show associations with multiple traits.

An additional challenge is that for every study that is added, the number of possible models of association doubles. In a large meta-analysis, it is computationally challenging to fully explore the possible subset space, and to conceptualize their prior probability because each model becomes more unlikely on its own. These challenges increase further if combinations of both positive and negative correlations are explicitly considered. It is possible that when the effect size estimates are precise (low *SE*), many of the subset models will have almost zero probability. In this scenario, the subset configuration space might be more efficiently explored by approaches such as Markov chain Monte Carlo (Smith & Roberts, 1993), or shotgun stochastic search (Hans et al., 2007). Here we have set parameters for the appropriate prior distributions. There is however considerable scope for fitting or estimating the parameters on the size and correlation of effects across models, and inferring the fraction of SNPs that derive from each of these models across the genome.

The method of genome-wide association study has become standard approach to gaining insights into the etiology of traits and disease—so common that it has be performed in an automated manner on thousands of traits in the UK Biobank (Canela-Xandri, Rawlik, & Tenesa, 2017; Howrigan, Abbott, Churchhouse, & Palmer, n.d.). As more biobanks are gathered, and as researchers start to peruse them for genetic associations, MetABF is a useful tool to determine cross-trait associations from GWAS. Its reliance on summary statistics means that access to the raw genetic data is not necessary. For a reasonable number of traits it can be applied genome-wide, overcoming some of the limitations of a traditional phenome-wide association study (PheWAS) (Cortes et al., 2017; Denny et al., 2010; Pendergrass et al., 2011). We have found the approach to aid in the assessment of which models of true associations are most consistent with the observed summary statistics at a variant.

## Supporting information

## 5 Acknowledgements

We thank the participants in the studies that generated the data for analysis within this manuscript. GM was supported by a Wellcome Trust grant (100956/Z/13/Z). CS was supported by a Wellcome Trust Career Development Fellowship grant (097364/Z/11/Z). MP was supported by grants from the Academy of Finland (288509, 294050, 312076). LJD is supported by the Wellcome Trust and the Royal Society (208750/Z/17/Z), and the Kennedy Trust for Rheumatology Research. CS is a shareholder and employee of Genomics PLC. GM is a director of and shareholder in Genomics PLC and a partner in Peptide Groove LLP. MP has provided consultancy services for Genomics plc. LJD has provided consultancy services for Genomics plc.

