## Supplementary material for "Bayesian meta-analysis across genome-wide association studies of diverse phenotypes"

### 6 Supplementary Material

#### 6.1 Statistical model

Logistic regression is a common approach to genome-wide association analysis of case-control studies. The regression approach allows for the inclusion of covariates such as age, sex, environment and ancestry. As genetic variants correlate with ancestry, and ancestry often correlates with case-control ascertainment, a failure to account for ancestry can inflate the evidence for association genome-wide (Cardon & Palmer, 2003; Price, Zaitlen, Reich, & Patterson, 2010). For a given SNP, let $g_i$ be the genotype of individual $i$ coded as the number of the reference alleles (0, 1, or 2) the individual is inferred to carry. Let $a_i$ also be the value of some covariate for individual $i$. If $p_i$ is the the probability that individual $i$ is a case, the logistic regression model with intercept $\beta_0$ and regression coefficients $\beta_g$ and $\beta_a$ can be written as:

$$\log\left(\frac{p_i}{1-p_i}\right) = \beta_0 + \beta_g g_i + \beta_a a_i. \tag{1}$$

The change in risk due to the reference allele is measured by the logarithm of the odds ratio (OR), equalling to $\beta_g g_i$, after adjusting for the covariates. The connection to the observed binary outcome $y_i \in \{0,1\}$, indicating control (0) or case (1) status, for a data set with $n$ individuals, is provided by the binomial likelihood function:

$$\mathcal{L}\left(\beta_0, \beta_g, \beta_a\right) = \prod_{i=1}^{n} p_i^{y_i}\left(1-p_i\right)^{1-y_i}. \tag{2}$$

There is no closed-form expression for the maximiser of this likelihood, and therefore it is usually estimated by an iterative algorithm. Assuming an additive model, where the log-odds of the disease increases (or decreases) linearly with the allelic dose, $\beta_g$ measures the effect of the reference allele on the risk of being a case. In what follows, we always consider the covariate-adjusted genetic effects and drop the $g$ subscript from the effect, i.e., $\beta = \beta_g$. Its standard error is estimated from the inverse of the observed information matrix at the maximum likelihood estimate (MLE) of the parameters (which measures the rate of change in the curvature of the likelihood surface). For quantitative traits, linear regression is a natural approach, in which case analytical results are available for the MLE of $\beta$ and the corresponding standard error. In large samples, the likelihood surface has a shape of a Gaussian distribution and therefore the MLE ($\hat{\beta}$) and its standard error ($SE_{\hat{\beta}}$) together are an accurate summary of the whole likelihood surface. They can be used for deriving several other quantities, such as the Wald test statistic:

$$z^2 = \frac{\left(\hat{\beta} - \beta\right)^2}{SE_{\hat{\beta}}^2},$$

which follows a $\chi_1^2$ distribution. The test statistic quantifies how far the observed $\hat{\beta}$ is from the true $\beta$. This test is used to assess the null hypothesis $\beta = 0$. Asymptotically, this test is equivalent to the likelihood ratio test (LRT), but computationally requires optimization for only one model (the alternative model), whereas the LRT requires optimization for two models (the null and the alternative models). Calculation of confidence intervals also relies on the summary statistics, for example, the 95% confidence interval is $\hat{\beta} \pm 1.96 \times SE_{\hat{\beta}}$.

Bayesian approaches are a natural way to include prior information about the magnitude of genetic effects on phenotypes, and the relationship of those effects across traits and studies. Bayesian approaches are built around Bayes' Theorem, which states that for any two events $A$ and $B$

$$\Pr(A|B) = \frac{\Pr(B|A)\Pr(A)}{\Pr(B)}. \tag{3}$$

The framework provides a natural way to compare two different models of the same data, $D$, e.g. the null model $M_0$ and some alternative model $M_1$, in terms of the posterior odds:

$$\underbrace{\frac{\mathrm{P}\left(M_1|D\right)}{\mathrm{P}\left(M_0|D\right)}}_{\text{Posterior odds}} = \underbrace{\frac{\mathrm{P}\left(D|M_1\right)}{\mathrm{P}\left(D|M_0\right)}}_{\text{Bayes factor}} \times \underbrace{\frac{\mathrm{P}\left(M_1\right)}{\mathrm{P}\left(M_0\right)}}_{\text{Prior odds}}. \tag{4}$$

The two terms that contribute to this ratio are the Bayes factor (BF) which is a weighting factor telling how the observed data updates the relative probabilities of the two models, and the prior odds describing the knowledge available before observing the data. Assume that $M_0$ and $M_1$ are parameterized by some sets of variables, $\theta_0$ and $\theta_1$, respectively. The Bayes factor requires computing marginal likelihoods for the models:

$$\mathrm{BF} = \frac{\mathrm{P}\left(D|M_1\right)}{\mathrm{P}\left(D|M_0\right)} = \frac{\int \mathrm{P}\left(D|\theta_1, M_1\right)\mathrm{P}\left(\theta_1|M_1\right)d\theta_1}{\int \mathrm{P}\left(D|\theta_0, M_0\right)\mathrm{P}\left(\theta_0|M_0\right)d\theta_0}. \tag{5}$$

The BF captures the information of the data on the relative likelihoods of the two models.

#### 6.2 Gaussian approximations

The integrals in Equation 5 are often difficult to evaluate, especially in the context of logistic regression discussed above. Because of this difficulty, we use an approximation developed by Wakefield specifically for GWAS (Wakefield, 2007, 2009). First we consider a single study. Let $\boldsymbol{y}$ be the vector of case-control labels, $\boldsymbol{X}$ matrix of covariates, $\boldsymbol{g}$ vector of genotypes, $\boldsymbol{\theta}$ vector of nuisance parameters, such as covariate effects, and $\beta$ the effect size of the SNP on the phenotype. Denote by $\ell$ the likelihood function of the logistic regression model and the prior distribution of $\boldsymbol{\theta}$ and $\beta$ by $p(\boldsymbol{\theta}, \beta)$.

Our goal is to approximate marginal likelihoods of the models, i.e., integrals of the type

$$\int \ell(\beta, \boldsymbol{\theta}|\boldsymbol{y}, \boldsymbol{X}, \boldsymbol{g}) p(\beta, \boldsymbol{\theta}) d\beta d\boldsymbol{\theta}.$$

Following Wakefield (Wakefield, 2007, 2009), we first approximate the likelihood function (up to a multiplicative constant $C$) by a multivariate normal density $f$ centred at the maximum likelihood estimate ($\hat{\beta}$,$\hat{\boldsymbol{\theta}}$) and having the covariance structure of the inverse of the observed information matrix $\hat{\boldsymbol{V}}_{\beta,\boldsymbol{\theta}}$:

$$\ell(\beta, \boldsymbol{\theta}; \boldsymbol{y}, \boldsymbol{X}, \boldsymbol{g}) \approx C \times f\left((\beta, \boldsymbol{\theta}) \mid (\hat{\beta}, \hat{\boldsymbol{\theta}}), \hat{\boldsymbol{V}}_{\beta,\boldsymbol{\theta}}\right). \tag{6}$$

We assume independent Gaussian prior distributions $p(\beta) = f(\beta|0, \Sigma_\beta)$ and $p(\theta) = f(\theta|0, \boldsymbol{\Sigma}_{\boldsymbol{\theta}})$ for parameters $\beta$ and $\boldsymbol{\theta}$, respectively. With these the marginal likelihood can be approximated as

$$\begin{aligned}\iint \ell(\boldsymbol{\theta}, \beta|\boldsymbol{y}, \boldsymbol{X}, \boldsymbol{g}) \quad &\times \quad p(\boldsymbol{\theta})p(\beta)\, d\beta d\boldsymbol{\theta} \\ &\approx \quad \iint C \times f\left((\beta, \boldsymbol{\theta})|(\hat{\beta}, \hat{\boldsymbol{\theta}}), \hat{V}_{\beta,\boldsymbol{\theta}}\right) f(\beta|0, \Sigma_\beta) f(\boldsymbol{\theta}|0, \boldsymbol{\Sigma}_{\boldsymbol{\theta}}) d\beta d\boldsymbol{\theta} \\ &= \quad C \times f\left((\hat{\beta}, \hat{\boldsymbol{\theta}})|(0,0), \hat{V}_{\beta,\boldsymbol{\theta}} + \mathrm{diag}(\Sigma_\beta, \boldsymbol{\Sigma}_{\boldsymbol{\theta}})\right) \\ &\approx \quad C \times f(\hat{\beta}|0, \hat{V}_\beta + \Sigma_\beta) f(\hat{\boldsymbol{\theta}}|0, \hat{V}_{\boldsymbol{\theta}} + \boldsymbol{\Sigma}_{\boldsymbol{\theta}}), \end{aligned} \tag{7}$$

where the last step assumes that we use a flat prior for nuisance parameters $\boldsymbol{\theta}$ where the diagonal elements of $\boldsymbol{\Sigma}_{\boldsymbol{\theta}}$ are very large allowing us to decouple $\beta$ and $\boldsymbol{\theta}$ even without assuming that they are independent in the likelihood.

Here we define our null model ($M_0$) to assume that there is no genetic effect, i.e. the prior on $\beta$ is a point mass at 0. The alternative model ($M_1$) assumes a prior distribution $\beta \sim \mathcal{N}(0, \sigma^2)$. For nuisance parameter $\boldsymbol{\theta}$ we assume the same flat prior in both models. The constant, $C$, and the normal density relating to the nuisance parameters, $f(\hat{\boldsymbol{\theta}}|\cdot)$ are not a function of $\beta$ and therefore cancel out from the top and bottom of BF. By combining formulas (5) and (7) we see that an approximate Bayes factor (ABF) between models $M_1$ and $M_0$ is :

$$\mathrm{ABF} = \frac{f(\hat{\beta}; 0, \hat{V}_\beta + \Sigma_\beta)}{f(\hat{\beta}; 0, \hat{V}_\beta)}. \tag{8}$$

For a single study, $\hat{V}_\beta = \mathrm{SE}^2_{\hat{\beta}}$ is the square of the observed standard error of $\hat{\beta}$ and $\Sigma_\beta = \sigma^2$ is the prior variance of the effect size. The ABF (8) generalizes to $n$ studies by assuming a multivariate normal prior on their effect sizes and a normal approximation to their joint likelihood. The covariance matrix for approximating the likelihood can be represented as $\hat{\boldsymbol{V}}_{\boldsymbol{\beta}} = \Delta(\mathrm{SE}_{\hat{\boldsymbol{\beta}}})\,\hat{\boldsymbol{R}}_{\boldsymbol{\beta}}\,\Delta(\mathrm{SE}_{\hat{\boldsymbol{\beta}}})$ where the correlation matrix of the effect size estimates across the studies, $\hat{\boldsymbol{R}}_{\boldsymbol{\beta}}$, can be estimated from genome-wide data as described in the main text and the diagonal matrix $\Delta(\mathrm{SE}_{\hat{\boldsymbol{\beta}}})$ contains the study specific standard errors of $\hat{\boldsymbol{\beta}}$ estimates. The covariance for the multivariate prior for $\boldsymbol{\beta}$ can be defined through the vector of study specific standard deviations $\boldsymbol{\sigma}$ and the correlation matrix $\boldsymbol{\rho}$ as $\boldsymbol{\Sigma}_{\boldsymbol{\beta}} = \Delta(\boldsymbol{\sigma})\,\boldsymbol{\rho}\,\Delta(\boldsymbol{\sigma})$.

#### 6.3 Simulations

We assessed whether sampling the effect size estimates from a Gaussian distribution yields the same distribution of $p$-values as simulating allele counts and performing the logistic regression. We conclude that when effect sizes are small or moderate, say OR of 1.1 as in Figure 6, the two simulation methods provide the same distribution, as long as the number of minor alleles in the sample is large enough (say several tens). In fact, Figure 6 is indistinguishable from the plot comparing the distributions resulting from these two methods under the null. However, when effect sizes are large, say OR of 1.5 as in Figure 7, one distribution tends to stochastically dominate the other.

**Figure 6:** Q-Q plots of the distribution of $p$-values for the different methods where OR $= 1.1$. The red line is $y = x$. The $x$-axis shows the distribution from simulations that used the logistic regression. The $y$-axis shows the distribution from sampling the effect size estimates directly from a Gaussian distribution.

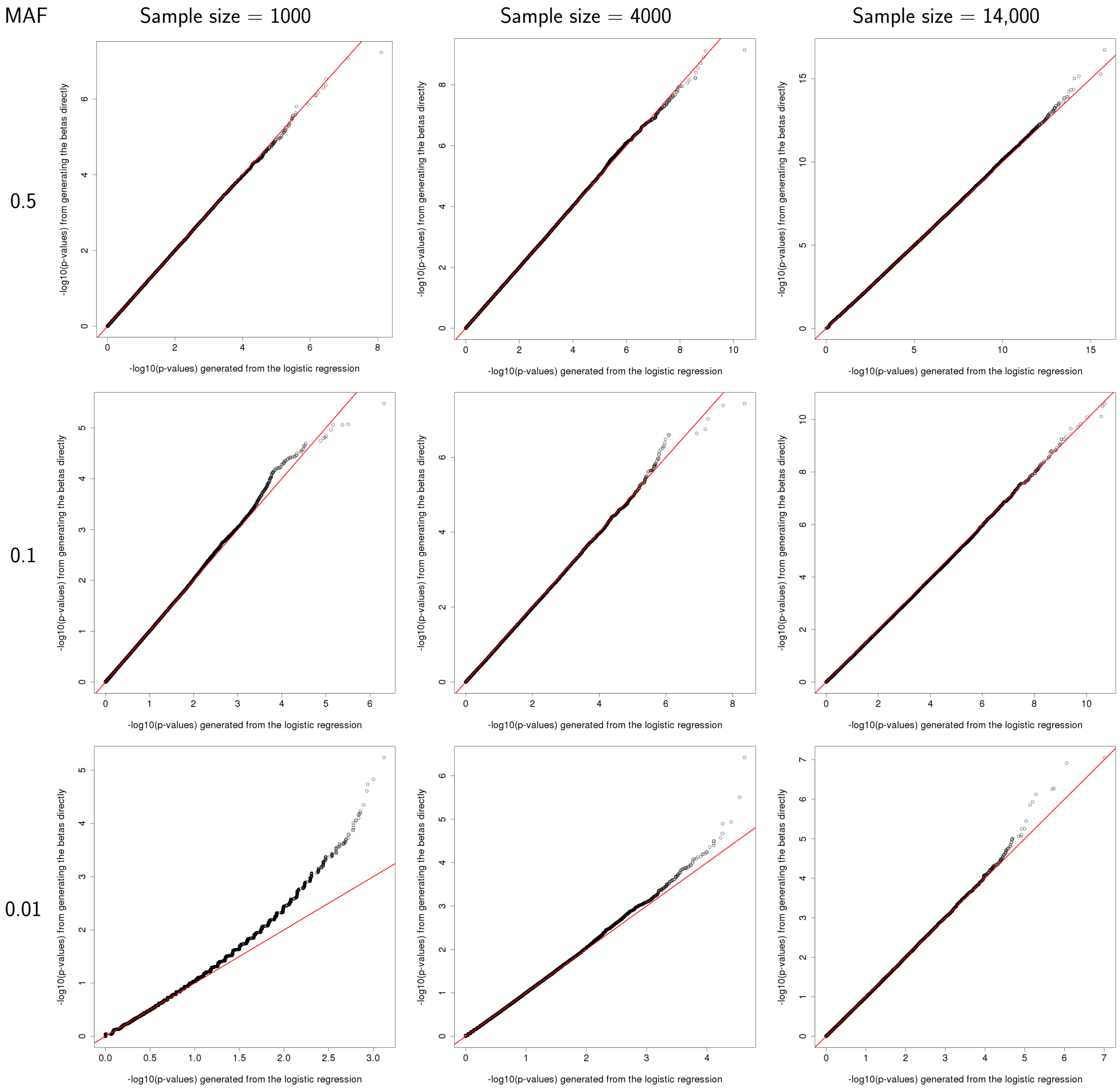

**Figure 7:** Q-Q plots on the distribution of $p$-values for the different methods where OR $= 1.5$. The red line is $y = x$. The $x$-axis shows the distribution from simulations that used the logistic regression. The $y$-axis shows the distribution from sampling the effect size estimates from a Gaussian distribution.

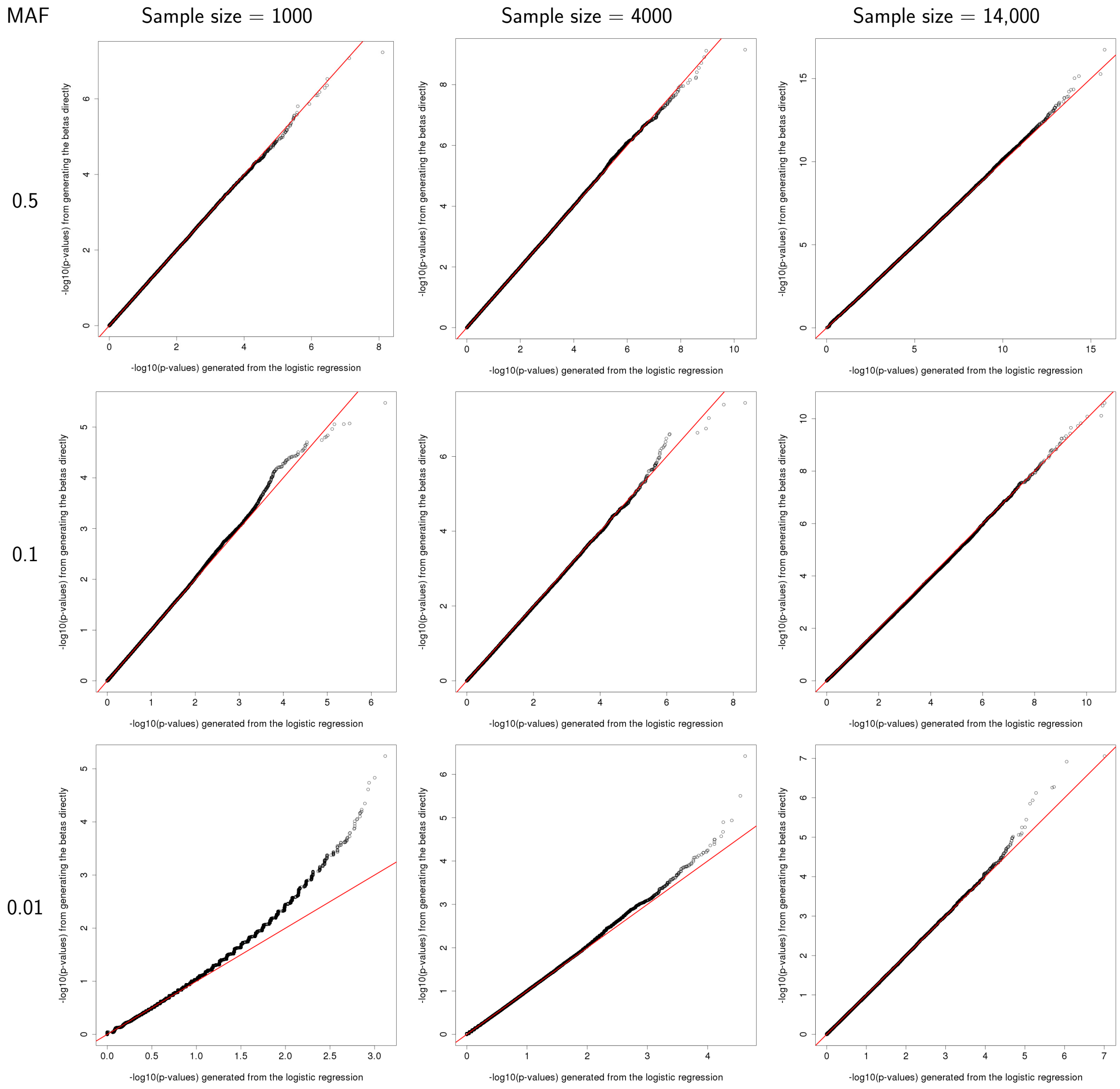

#### 6.4 Power analyses

| | | Number of phenotypes associated at $\sigma_{gen}$=1e−05 | | | | | Number of phenotypes associated at $\sigma_{gen}$=0.1 | | | | | Number of phenotypes associated at $\sigma_{gen}$=0.15 | | | | |
|---|---|---|---|---|---|---|---|---|---|---|---|---|---|---|---|---|
| | | 1 | 2 | 3 | 4 | 5 | 1 | 2 | 3 | 4 | 5 | 1 | 2 | 3 | 4 | 5 |
| | Fixed effects | 0.019 | 0.029 | 0.042 | 0.055 | 0.068 | 0.233 | 0.454 | 0.587 | 0.67 | 0.727 | 0.39 | 0.605 | 0.711 | 0.773 | 0.813 |
| | Random effects | 0.019 | 0.029 | 0.042 | 0.055 | 0.068 | 0.233 | 0.454 | 0.587 | 0.67 | 0.727 | 0.39 | 0.604 | 0.71 | 0.771 | 0.811 |
| | Fisher's method | 0.031 | 0.06 | 0.095 | 0.136 | 0.181 | 0.44 | 0.701 | 0.846 | 0.923 | 0.963 | 0.594 | 0.841 | 0.94 | 0.979 | 0.992 |
| $\rho_{an}$=0 | corr BF | 0.033 | 0.063 | 0.099 | 0.139 | 0.183 | 0.458 | 0.713 | 0.852 | 0.926 | 0.963 | 0.609 | 0.849 | 0.943 | 0.979 | 0.993 |
| | mean BF | 0.037 | 0.066 | 0.098 | 0.132 | 0.167 | 0.485 | 0.725 | 0.852 | 0.921 | 0.958 | 0.631 | 0.856 | 0.943 | 0.978 | 0.991 |
| $\rho_{an}$=0.5 | corr BF | 0.033 | 0.063 | 0.099 | 0.139 | 0.183 | 0.458 | 0.714 | 0.853 | 0.926 | 0.963 | 0.609 | 0.849 | 0.943 | 0.979 | 0.993 |
| | mean BF | 0.037 | 0.067 | 0.098 | 0.133 | 0.168 | 0.485 | 0.725 | 0.853 | 0.922 | 0.959 | 0.631 | 0.856 | 0.943 | 0.978 | 0.991 |
| $\rho_{an}$=0.96 | corr BF | 0.033 | 0.062 | 0.096 | 0.136 | 0.179 | 0.455 | 0.711 | 0.851 | 0.925 | 0.963 | 0.606 | 0.847 | 0.942 | 0.979 | 0.992 |
| | mean BF | 0.036 | 0.066 | 0.098 | 0.134 | 0.171 | 0.479 | 0.723 | 0.854 | 0.923 | 0.96 | 0.626 | 0.854 | 0.943 | 0.978 | 0.992 |

**Figure 8:** Heatmap and table of statistical power for $\alpha = 0.01$ comparing the ABF approach using different parameters against frequentist meta-analysis: inverse-variance weighted fixed effects, random effects calculated in R using the DerSimonian and Laird method (DerSimonian & Laird, 1986), and Fisher's method. We show the true positive rate at this level for data where 1-5 phenotypes were affected and where the effect sizes were generated using different values of the prior $\sigma_{gen}$. All data were generated using a prior $\rho_{gen} = 0.5$ and analyzed using a prior value of $\sigma = 0.2$. The size of the study was 7000 cases and 7000 controls. The minor allele frequency was 0.5. The prior weight on all models was equal. A null dataset of 100,000,000 SNPs was created for which the all correlated and mean Bayes factors across all subset models (for each value of $\rho_{an}$) were calculated. This left us with a null distribution for each Bayes factor approach, allowing us to compute empirical $p$-values.

| | Number of phenotypes associated at $\rho_{gen}$=0 | | | | | Number of phenotypes associated at $\rho_{gen}$=0.5 | | | | | Number of phenotypes associated at $\rho_{gen}$=0.96 | | | | | Number of phenotypes associated at $\rho_{gen}$=1 | | | | |
|---|---|---|---|---|---|---|---|---|---|---|---|---|---|---|---|---|---|---|---|---|
| | 1 | 2 | 3 | 4 | 5 | 1 | 2 | 3 | 4 | 5 | 1 | 2 | 3 | 4 | 5 | 1 | 2 | 3 | 4 | 5 |
| Fixed effects | 0.507 | 0.629 | 0.691 | 0.73 | 0.757 | 0.507 | 0.693 | 0.779 | 0.827 | 0.858 | 0.507 | 0.731 | 0.817 | 0.862 | 0.889 | 0.506 | 0.733 | 0.819 | 0.864 | 0.891 |
| Random effects | 0.507 | 0.617 | 0.662 | 0.682 | 0.689 | 0.507 | 0.691 | 0.774 | 0.82 | 0.847 | 0.507 | 0.731 | 0.817 | 0.862 | 0.889 | 0.506 | 0.733 | 0.819 | 0.864 | 0.891 |
| Fisher's method | 0.686 | 0.915 | 0.979 | 0.995 | 0.999 | 0.686 | 0.904 | 0.972 | 0.992 | 0.998 | 0.686 | 0.818 | 0.885 | 0.925 | 0.951 | 0.686 | 0.79 | 0.84 | 0.871 | 0.893 |
| $\rho_{an}$=0 corr BF | 0.698 | 0.92 | 0.98 | 0.996 | 0.999 | 0.698 | 0.909 | 0.973 | 0.992 | 0.998 | 0.698 | 0.825 | 0.889 | 0.927 | 0.952 | 0.698 | 0.795 | 0.842 | 0.872 | 0.893 |
| $\rho_{an}$=0 mean BF | 0.716 | 0.923 | 0.98 | 0.995 | 0.999 | 0.716 | 0.914 | 0.973 | 0.992 | 0.998 | 0.716 | 0.83 | 0.888 | 0.923 | 0.946 | 0.716 | 0.797 | 0.838 | 0.865 | 0.885 |
| $\rho_{an}$=0.5 corr BF | 0.698 | 0.92 | 0.98 | 0.995 | 0.999 | 0.698 | 0.909 | 0.973 | 0.992 | 0.998 | 0.698 | 0.825 | 0.889 | 0.927 | 0.952 | 0.698 | 0.795 | 0.842 | 0.873 | 0.894 |
| $\rho_{an}$=0.5 mean BF | 0.716 | 0.924 | 0.98 | 0.995 | 0.999 | 0.716 | 0.914 | 0.973 | 0.992 | 0.998 | 0.716 | 0.831 | 0.888 | 0.924 | 0.947 | 0.716 | 0.797 | 0.839 | 0.866 | 0.886 |
| $\rho_{an}$=0.96 corr BF | 0.696 | 0.918 | 0.98 | 0.995 | 0.999 | 0.696 | 0.908 | 0.973 | 0.992 | 0.998 | 0.696 | 0.825 | 0.89 | 0.928 | 0.952 | 0.696 | 0.798 | 0.847 | 0.878 | 0.9 |
| $\rho_{an}$=0.96 mean BF | 0.712 | 0.922 | 0.98 | 0.995 | 0.999 | 0.712 | 0.913 | 0.973 | 0.992 | 0.998 | 0.712 | 0.832 | 0.891 | 0.927 | 0.95 | 0.712 | 0.801 | 0.846 | 0.875 | 0.895 |

**Figure 9:** Heatmap and table of statistical power for $\alpha = 0.01$ comparing the ABF approach using different parameters against frequentist meta-analysis: inverse-variance weighted fixed effects, random effects calculated in R using the DerSimonian and Laird method (DerSimonian & Laird, 1986), and Fisher's method. We show the true positive rate at this level for data where 1-5 phenotypes were affected and where the effect sizes were generated using different values of the correlation coefficient of $\rho_{gen}$. All data were generated using a prior $\sigma_{gen} = 0.2$ and analyzed using a prior $\sigma = 0.2$. The size of the study was 7000 cases and 7000 controls and the minor allele frequency was 0.5. The prior weight on all models was equal. A null dataset of 100,000,000 SNPs was created for which the all correlated and mean Bayes factors across all subset models (for each value of $\rho_{an}$) were calculated). This left us with a null distribution for each Bayes factor approach, allowing us to compute empirical $p$-values.

#### 6.5 $\boldsymbol{P}$-values and null distributions

The multivariate Gaussian approach described here in a Bayesian setting can be viewed as either a model comparison test, or a simple test statistic. Like any test statistic, the ABF has a null distribution, which depends on the covariance between the studies, $\boldsymbol{V}_{\hat{\boldsymbol{\beta}}}$, and the prior used in the analysis, $\boldsymbol{\Sigma}$. As discussed in the analysis of the WTCCC2 data (Section 3.4), for a given $\boldsymbol{V}_{\hat{\boldsymbol{\beta}}}$ and $\boldsymbol{\Sigma}$, it is fast to generate data under the null and calculate their corresponding ABFs to get the null distribution of the test statistic. Comparing the ABFs calculated from the data to this null distribution allows for the calculation of $p$-values.

The ABF in formula (4) depends on $\hat{\beta}$ only through the following quadratic form in a monotonic manner,

$$Q\left(\hat{\boldsymbol{\beta}}; \boldsymbol{V}_{\hat{\boldsymbol{\beta}}}, \boldsymbol{\Sigma}\right) = \hat{\boldsymbol{\beta}}^{\top}\left(\boldsymbol{V}_{\hat{\boldsymbol{\beta}}}^{-1} - (\boldsymbol{V}_{\hat{\boldsymbol{\beta}}} + \boldsymbol{\Sigma})^{-1}\right)\hat{\boldsymbol{\beta}}. \tag{9}$$

Under the null model $M_0$: $\hat{\boldsymbol{\beta}} \sim \text{MVN}(\mathbf{0}, \boldsymbol{V}_{\hat{\boldsymbol{\beta}}})$ and hence $Q(\hat{\boldsymbol{\beta}}; \boldsymbol{V}_{\hat{\boldsymbol{\beta}}}, \boldsymbol{\Sigma}) \sim \sum_{i=1}^{n} d_i \chi_i^2$, where $\chi_i^2$ are an independent sample from $\chi_1^2$ distribution (chi-square with one degree of freedom), and $d_i$ are the eigenvalues of matrix $I - (\boldsymbol{V}_{\hat{\boldsymbol{\beta}}} + \boldsymbol{\Sigma})^{-1} \boldsymbol{V}_{\hat{\boldsymbol{\beta}}}$. The distribution function for a mixture of chi-squares can be numerically evaluated, for example by the R-package 'CompQuadForm' (Duchesne & de Micheaux, 2010). This allows us to compute $p$-values under the null model directly by using ABF as the test statistic, permitting a full range of frequentist meta-analysis tests (fixed effects, correlated effects, through to independent effects), which can cope with arbitrary structure in the covariance of the noise between studies, and including information on the size of plausible effects.

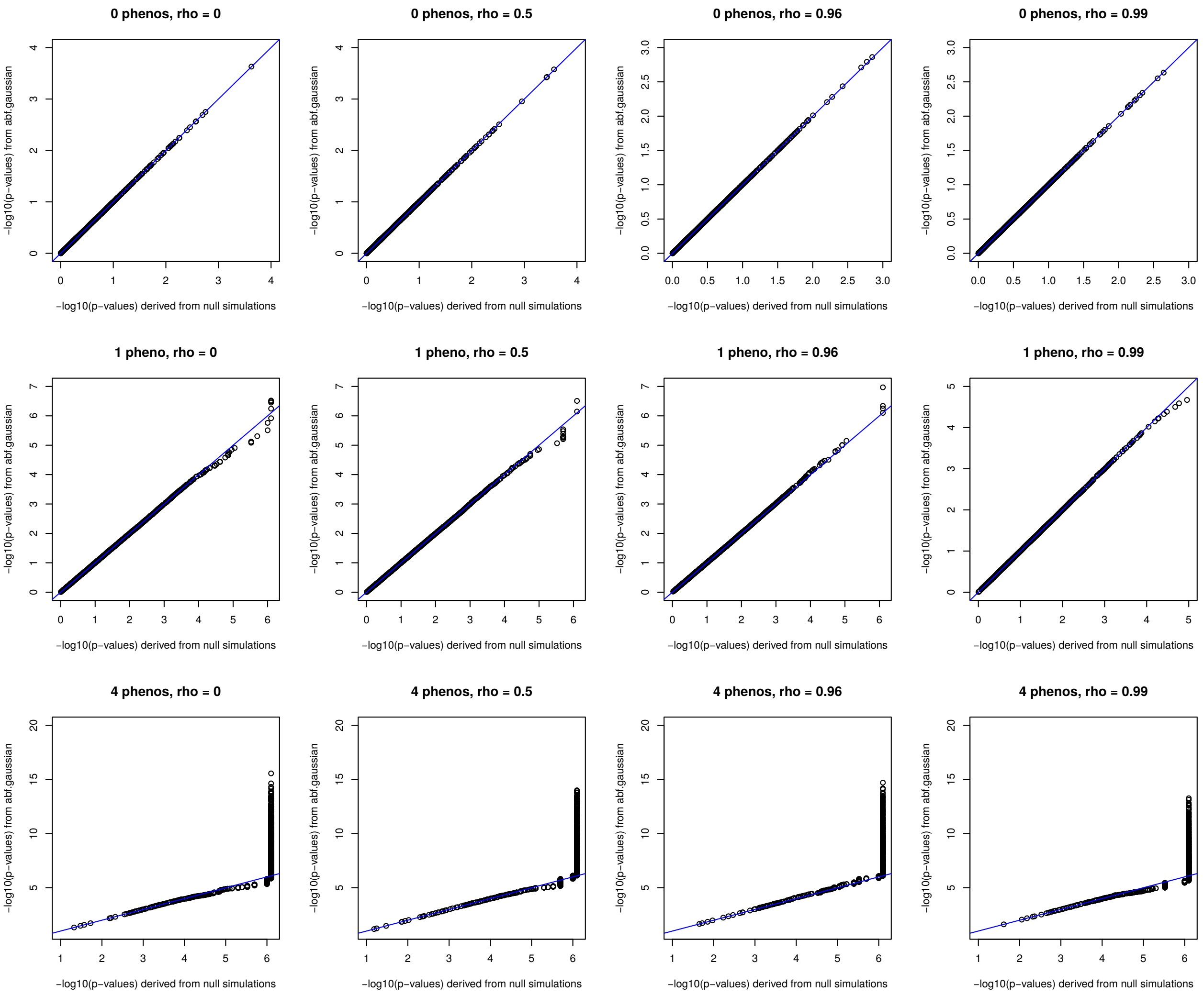


**Figure 10:** Plots showing $p$-values calculated using the analytic method ("abf.gaussian"), and those using a null distribution of one million SNPs. The blue line indicates $y = x$. Further simulated data were generated with true effects in 0, 1, and 4 out of a possible 7 studies ("phenos"). Where true associations existed, the odds ratio was 1.1. Data were analyzed with $\sigma = 0.2$ and with different values of $\rho$ ("rho"). We see good concordance between the analytic $p$-values and those from the null distribution until $p \approx 10^{-5}$, which is where the cut-offs determined by the null distribution become less reliable, due to few SNPs generating ABFs this extreme under the null.

#### 6.6 Model selection – sensitivity and specificity

When performing the MetABF calculation for every possible subset of association, it is natural to choose the model with the highest ABF as the most likely model. Here we examine the accuracy of this approach to model selection in simulated data. Summary statistics were simulated for six million SNPs across five studies. True effects were drawn randomly from a normal distribution with standard deviation $\sigma_{gen} = 0.2$ and correlations between the studies of $\rho_{gen} = \{0, 0.5, \pm 0.5, 1\}$, where $\rho = \pm 0.5$ uses the prior matrix equation 10 below:

$$\begin{bmatrix} 1 & -0.5 & 0.5 & -0.5 & 0.5 \\ -0.5 & 1 & -0.5 & 0.5 & -0.5 \\ 0.5 & -0.5 & 1 & -0.5 & 0.5 \\ -0.5 & 0.5 & -0.5 & 1 & -0.5 \\ 0.5 & -0.5 & 0.5 & -0.5 & 1 \end{bmatrix} \tag{10}$$

MetABF analysis was done using $\sigma = 0.2$ and different values of $\rho$. In each analysis, the model with the highest ABF was compared to the true underlying model of association. We report the sensitivity (proportion of times the chosen model declares a true association in a given study when the study has a true association) and the specificity (proportion of times the chosen model declares no association in a given study when there is no true association in the study) in Tables 3-6 below.

**Table 3:** The sensitivity and specificity resulting from selecting the model with the highest Bayes factor using different values of the prior $\rho$ in the analysis and $\sigma_i = 0.2$. The data were generated for five traits by drawing the effect size from a normal distribution, using a correlation of $\rho_{gen} = 0$ for all pairs of traits and $\sigma_{gen} = 0.2$ for all traits.

| Model | $\rho = 0$ | $\rho = 0.5$ | $\rho = 0.75$ | $\rho = 0.96$ | $\rho = 1$ |
|---|---|---|---|---|---|
| Sensitivity | 0.806 | 0.806 | 0.800 | 0.694 | 0.528 |
| Specificity | 0.962 | 0.962 | 0.959 | 0.959 | 0.975 |

We see from the tables that the analysis with the parameter $\rho = 0$ gives surprisingly consistent results—from the dataset generated using $\rho = 0$ for all pairs of correlations to the

**Table 4:** The sensitivity and specificity resulting from selecting the model with the highest Bayes factor using different values of the prior $\rho$ in the analysis and $\sigma_i = 0.2$. The data were generated for five traits by drawing the effect size from a normal distribution, using a correlation of $\rho_{gen} = 0.5$ for all pairs of traits and $\sigma_{gen} = 0.2$ for all traits.

| Model | $\rho = 0$ | $\rho = 0.5$ | $\rho = 0.75$ | $\rho = 0.96$ | $\rho = 1$ |
|---|---|---|---|---|---|
| Sensitivity | 0.806 | 0.807 | 0.803 | 0.732 | 0.623 |
| Specificity | 0.962 | 0.964 | 0.964 | 0.966 | 0.973 |

**Table 5:** The sensitivity and specificity resulting from selecting the model with the highest Bayes factor using different values of the prior $\rho$ in the analysis and $\sigma_i = 0.2$. The data were generated for five traits by drawing the effect size from a normal distribution, using a correlation of $\rho_{gen} = 1$ for all pairs of traits and $\sigma_{gen} = 0.2$ for all traits.

| Model | $\rho = 0$ | $\rho = 0.5$ | $\rho = 0.75$ | $\rho = 0.96$ | $\rho = 1$ |
|---|---|---|---|---|---|
| Sensitivity | 0.806 | 0.812 | 0.820 | 0.840 | 0.846 |
| Specificity | 0.962 | 0.965 | 0.965 | 0.965 | 0.959 |

**Table 6:** The sensitivity and specificity resulting from selecting the model with the highest Bayes factor using different (positive) values of the prior $\rho$ in the analysis and $\sigma_i = 0.2$. The data were generated for five traits by drawing the effect size from a normal distribution, using a correlation of $\rho_{gen} = \pm 0.5$ for all pairs of traits and $\sigma_{gen} = 0.2$ for all traits.

| Model | $\rho = 0$ | $\rho = 0.5$ | $\rho = 0.75$ | $\rho = 0.96$ | $\rho = 1$ |
|---|---|---|---|---|---|
| Sensitivity | 0.806 | 0.807 | 0.805 | 0.710 | 0.528 |
| Specificity | 0.962 | 0.961 | 0.957 | 0.955 | 0.974 |

one where $\rho = \pm 0.5$. This is in part due to rounding—keeping a few extra digits shows the differences in the performance of analyses with this parameter. We also see that specificity is always greater than 0.95, which means that we do not call false positives more than 5% of the time. That said, there is quite a range in sensitivities, which drop to almost 0.5 when $\rho = 1$ and the data were generated using a different value. In fact, these tables indicate that performing the analysis with $\rho = 0.96$ will yield results that are at least as accurate as running it with $\rho = 1$, including when the data were generated by $\rho = 1$.

We also notice the surprising result that the accuracy of this approach to model selection does not appear to be harmed even when the sign of the correlation coefficient is incorrect. Table 6—where the values of $\rho$ were $\pm 0.5$, according to matrix 10—is entirely consistent with the others, despite the fact that we would expect lower values across all analyses due to the true underlying negative correlations between some of the studies being assumed to be positively correlated in the analysis. It may be because if the sign of $\rho$ is incorrect, then it is equally incorrect in all models of association and thus does not affect their final rankings.

Finally, we see that the main effect of choosing $\rho$ wisely is on power to detect a true effect. This is somewhat comforting—if $\rho$ is not chosen well, then the model corresponding to the highest Bayes factor will call extra false negatives, but it should not produce excessive false positives.

#### 6.7 WTCCC2 studies

Processed GWAS summary statistics will be available from the European Genotype Archive and information on how to apply for access to the data is given found here :
https://www.wtccc.org.uk/info/access_to_data_samples.html.

##### 6.7.1 Preprocessing steps

We obtained summary statistics for each SNP, its chromosome and base pair position (on NCBI human genome build 36), rsid, alleles (coded "allele A" and "allele B"), frequency of the B allele in controls, the estimated effect size of the SNP on the trait, standard error of the estimated effect size, and the $p$-value. Associated with each file was a list of SNP inclusions or exclusions from the original studies. These filters were applied to ensure that only the SNPs that passed quality control in the original studies were included in the meta-analysis.

In order to be able to perform a meaningful meta-analysis, we aimed to ensure that the A and B alleles of each SNP were the same in all studies (so that the direction of effect is measure consistently). We did this by aligning our alleles to the forward strand and checking that the reported allele frequencies matched those in the 1000 Genomes Project. We describe these steps below.

The first step was to check the internal consistency between the effect size estimates and their standard errors and the reported $p$-values. In Figure 11, we see that the $p$-values calculated from the effect size estimates and standard errors are positively correlated ($\rho \approx 0.99997$) with those reported in the summary statistic tables and that while these values do not always match perfectly, the results do appear to be consistent with one another. This suggests that there have not been major errors in the tables of summary statistics for each study and that the effect size estimates and $p$-values are generally being reported correctly.

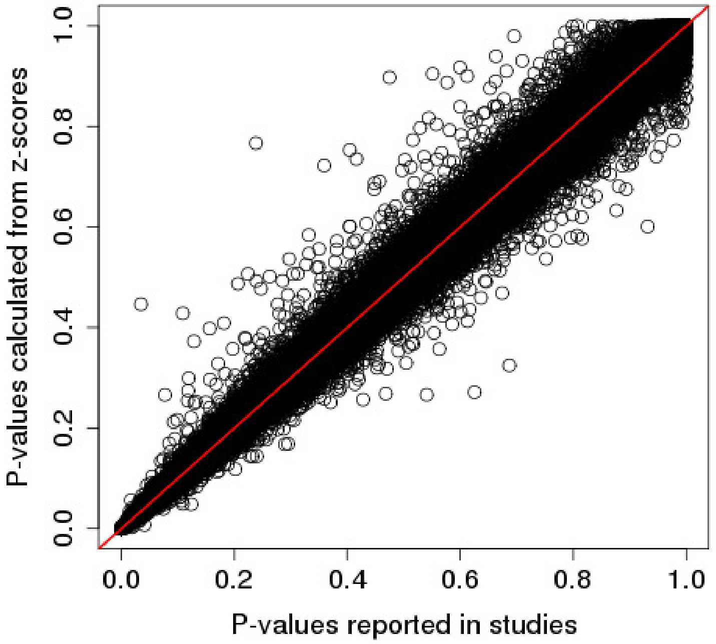


**Figure 11:** Comparison of the $p$-values reported in the WTCCC2 tables of summary statistics and the $p$-values calculated from the $z$-scores (calculated from the reported effect size estimates and standard errors) using a $\chi^2_1$ test. The red line is $y = x$.

The next step was to compare the rsids and positions of each SNP to those in a publicly available set of curated genotyping array manifest which contained information about the strand on which each allele is assayed (Rayner, 2017). For each study, there were a small number of anomalous SNPs whose positions were not found in the manifests and were thus removed. The next step was to align all SNPs to the forward (+) strand. In the case of SNPs that were aligned to the backward (-) strand the alleles were complemented so that they were on the forward strand. We then aligned the alleles with the reference sequence by swapping Allele A and Allele B around, subtracting the stated allele frequency from 1, and changing the sign of the estimated effect size as required.

We then compared each file to the 1000 Genomes Phase 3 data. The SNPs in 1000 Genomes are all aligned to the forward strand—the "ref" allele was expected to correspond to our allele A and "alt" was expected to correspond to our allele B, so this step was taken to ensure that the alleles were correct. SNPs whose alleles did not match 1000 Genomes were removed from the data. Finally, we used the allele frequencies in the study and the reference populations to calculate Wright's $F_{ST}$ between them and removed any SNPs where $F_{ST} \geq 0.1$. As a final check, we plotted the allele frequencies of the SNPs listed in the original files for each study before any alignment processing was done and compared that to the corresponding plot after the alignment. We show these plots for the psoriasis study (PS) in Figure ??.

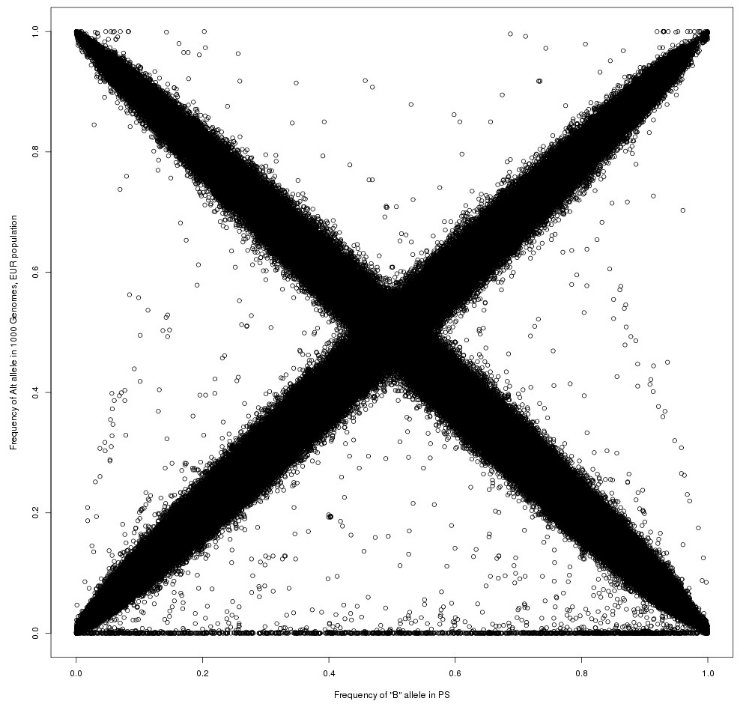


**Figure 12:** Allele frequencies of the alt allele from the 1000 Genomes EUR population plotted against the allele frequencies of the B allele in the psoriasis study, before alignment and filtering.

We were satisfied with our pipeline for other phenotypes except for PR, which we

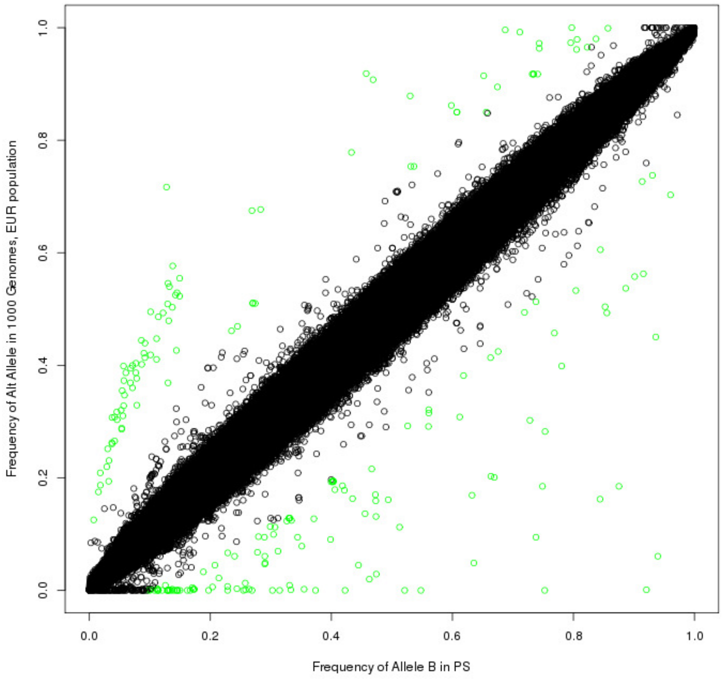


**Figure 13:** Allele frequencies of the alt allele from the 1000 Genomes EUR population plotted against the allele frequencies of the B allele in the psoriasis study, after alignment and filtering. Green circles represent the SNPs that were filtered out due to $F_{ST} \geq 0.1$.

subsequently excluded from our analyses. This left us with 20 individual cohorts across 14 studies. We ran our meta-analysis method on the 2,058,911 SNPs for which we had data—effect size estimates and their associated standard errors—in at least two cohorts.

##### 6.7.2 Data description

**Table 7:** Summary of the data available from the Wellcome Trust Case Control Consortium Study 2. We list the phenotypes, the abbreviation we use for them ("Abbr."), the genotyping chips (cases/controls), the reference panel to which they were imputed if applicable, and the reference to the published paper in the bibliography.

| **Phenotype** | **Abbr.** | **Genotyping chip (case/control)** | **Reference panel** | **Reference** |
|---|---|---|---|---|
| Ulcerative colitis | UC | Affymetrix 6.0 SNP array | Not imputed | Barrett et al. (2009) |
| Psoriasis | PS | Illumina 660W-Quad array/Illumina Human 1.2M-Duo array | Not imputed | Genetic Analysis of Psoriasis Consortium and The Wellcome Trust Case Control Consortium 2 (2010) |
| Parkinson's disease | PD | Illumina 660W-Quad array/Illumina Human 1.2M-Duo array | Not imputed | The UK Parkinson's Disease Consortium and The Wellcome Trust Case Control Consortium 2 et al. (2011) |
| Glycemic response to metformin in type 2 diabetes | PR | Affymetrix 6.0 SNP array | Not imputed | Zhou et al. (2011) |
| Ankylosing spondylitis | AS | Illumina 660W-Quad array/Illumina Human 1.2M-Duo array | HapMap2 CEU, HapMap3 CEU | Evans et al. (2011) |
| Multiple sclerosis | MS | Illumina 660W-Quad array/Illumina Human 1.2M-Duo array | Not imputed | Sawcer et al. (2011) |
| Ischemic stroke | IS | Illumina 660W-Quad array/Illumina Human 1.2M-Duo array | Not imputed | Bellenguez et al. (2012) |

*(Continued on next page)*

*(continued from previous page)*

| Phenotype | Abbr. | Genotyping chip (case/control) | Reference panel | Reference |
|---|---|---|---|---|
| Barrett's esophagus | BO | Illumina 660W-Quad array/Illumina Human 1.2M-Duo array | Not imputed | Su et al. (2012) |
| Schizophrenia, schizoaffective disorder, and schizophreniform disorder | SP | Affymetrix 6.0 SNP array | Not imputed | Irish Schizophrenia Genomics Consortium and Wellcome Trust Case Control Consortium 2 (2012) |
| Visceral leishmaniasis | VL | Illumina 660W-Quad array | Not imputed | Fakiola et al. (2013) |
| Intra-ocular pressure (glaucoma endophenotype) | GL | Illumina 660W-Quad array | 1000 Genomes Pilot CEU haplotypes | The Blue Mountains Eye Study (BMES) and The Wellcome Trust Case Control Consortium 2 (WTCCC2) et al. (2013) |
| Psychosis | PE | Affymetrix 6.0 SNP array | Not imputed | Psychosis Endophenotypes International Consortium and Wellcome Trust Case-Control Consortium 2 (2014) |
| Reading and mathematical performance at age 12 | RM | Affymetrix 6.0 SNP array | HapMap2 CEU, HapMap3 CEU | Davis et al. (2014) |
| Bacteremia susceptibility | BS | Affymetrix 6.0 SNP array | 1000 Genomes Phase 1 | Rautanen et al. (2016) |

**Table 8:** Description of each study (subdivided into cohorts, if necessary), with the total numbers of cases and controls, number of controls from the WTCCC2 pool of shared controls, the population from which the samples were drawn, and any miscellaneous comments. Separate cohorts are appended. For instance, the visceral leishmaniasis cohort from India is labelled VL India. For studies with multiple cohorts, we give the total numbers across all studies. These rows are italicized. For studies investigating a quantitative trait, the total number of participants is listed in the cases column.

| Study | Cases | Controls | WTCCC2 Controls | Population | Notes |
|---|---|---|---|---|---|
| UC | 2361 | 5417 | 5417 | European | |
| PS | 2178 | 5175 | 5175 | European | |
| PD | 1705 | 5175 | 5175 | European | |
| PR | 1024 | | | European | |
| AS | 1787 | 4800 | 4800 | European | |
| MS UK | 1854 | 5175 | 5175 | European | |
| MS nonUK | 7918 | 12201 | 0 | European | |
| *MS* | *9772* | *17376* | *5175* | *European* | |
| IS TOAST 1 | 844 | 5972 | 5175 | European | Large vessel stroke cohort |
| IS TOAST 2 | 580 | 5972 | 5175 | European | Small vessel stroke cohort |
| IS TOAST 3 | 790 | 5972 | 5175 | European | Cardioembolic stroke cohort |
| *IS* | *3548* | *5972* | *5175* | *European* | *Includes samples not in TOAST cohorts.* |
| BO | 1852 | 5172 | 5172 | European | |
| SP | 1606 | 1794 | 0 | European (Irish) | |
| VL India | 989 | 1089 | 0 | Indian | |
| VL Brazil | 357 | 1613 | 0 | Brazilian | Cases and controls drawn from 308 families. |
| *VL* | *1346* | *2702* | *0* | *Indian and Brazilian* | |
| GL | 2302 | | | European | |

*(Continued on next page)*

*(continued from previous page)*

| Study | Cases | Controls | WTCCC2 Controls | Population | Notes |
|---|---|---|---|---|---|
| PE | 1239 | 3596 | 0 | European | 857 of the controls are unaffected relatives of cases. |
| RM | 2794 | | | European | |
| BS pneumococcal | 429 | 2677 | 0 | Kenyan | |
| BS overall | 1536 | 2677 | 0 | Kenyan | |

##### 6.7.3 WTCCC2 Q-Q plot of ABF distribution

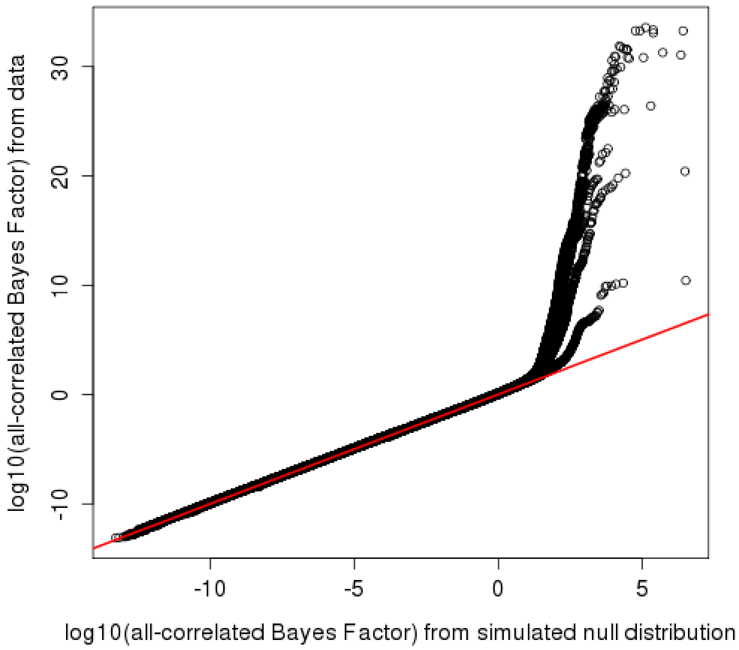


**Figure 14:** Q-Q plot of the null distribution of approximate Bayes factors against the ABFs calculated from the data using each of the 12 possible correlation matrices and a prior $\sigma = 0.2$. The cryptic relatedness among the studies was accounted for by using the covariance matrix calculated from the data with the genome-wide significant signals removed.

##### 6.7.4 Results from the individual GWAS

**Table 9:** Non-HLA SNPs that had been previously implicated before each of the WTCCC2 genome-wide association studies. We provide the SNP rsid, the disease with which it was previously thought to be associated, the $p$-value reported in the WTCCC2 study (discovery cohort), and the mean ABF across our analyses. Note that some SNPs appear more than once due to their association with multiple traits. SNPs for which there was summary statistic data for only one trait could not have a MetABF analysis performed, and have in NA the "mean ABF" column.

| SNP | Disease | $P$-value in study | mean ABF |
|---|---|---|---|
| rs11209026 | AS | $1.9 \times 10^{-6}$ | 14.51 |
| rs10865331 | AS | $2.9 \times 10^{-12}$ | 14.26 |

*(Continued on next page)*

*(continued from previous page)*

| **SNP** | **Disease** | ***p*-value in study** | **mean ABF ($\log_{10}$)** |
|---|---|---|---|
| rs30187 | AS | $4.3 \times 10^{-13}$ | 17.26 |
| rs4349859 | AS | $1 \times 10^{-200}$ | NA |
| rs378108 | AS | $1.5 \times 10^{-7}$ | 7.07 |
| rs11209026 | PS | $7.13 \times 10^{-7}$ | 14.51 |
| rs4112788 | PS | $3.32 \times 10^{-10}$ | 3.82 |
| rs3213094 | PS | $4.93 \times 10^{-11}$ | 5.69 |
| rs610604 | PS | $6.65 \times 10^{-7}$ | 2.70 |
| rs2066808 | PS | $2.49 \times 10^{-7}$ | 1.02 |
| rs4648356 | MS | $3.1 \times 10^{-14}$ | 7.88 |
| rs11810217 | MS | $6.5 \times 10^{-12}$ | 6.00 |
| rs1335532 | MS | $2 \times 10^{-9}$ | 6.23 |
| rs1323292 | MS | $8.8 \times 10^{-7}$ | 2.76 |
| rs7522462 | MS | $9.2 \times 10^{-7}$ | 8.42 |
| rs2293370 | MS | $1.1 \times 10^{-9}$ | 3.32 |
| rs4613763 | MS | $6.9 \times 10^{-14}$ | 9.10 |
| rs1520333 | MS | $6.1 \times 10^{-7}$ | 0.60 |
| rs3118470 | MS | $2 \times 10^{-9}$ | 3.17 |
| rs650258 | MS | $1.7 \times 10^{-7}$ | 3.15 |
| rs1800693 | MS | $1.8 \times 10^{-10}$ | 8.33 |
| rs12368653 | MS | $2 \times 10^{-7}$ | 2.36 |
| rs7200786 | MS | $6.3 \times 10^{-14}$ | 6.83 |
| rs13333054 | MS | $7 \times 10^{-8}$ | 2.26 |
| rs9891119 | MS | $4.6 \times 10^{-7}$ | 4.24 |

*(Continued on next page)*

(continued from previous page)

| SNP | Disease | $p$-value in study | mean ABF ($\log_{10}$) |
|---|---|---|---|
| rs180515 | MS | $1.4 \times 10^{-7}$ | 2.24 |
| rs6062314 | MS | $8.3 \times 10^{-7}$ | 3.04 |
| rs6426833 | UC | $2.1 \times 10^{-11}$ | 4.90 |
| rs11209026 | UC | $3 \times 10^{-10}$ | 14.51 |
| rs3024493 | UC | $8 \times 10^{-8}$ | 2.98 |
| rs9858542 | UC | $7 \times 10^{-7}$ | 4.02 |
| rs6584283 | UC | $1.7 \times 10^{-7}$ | 3.12 |
| | IS TOAST | | |
| rs1906599 | 3 | $3.45 \times 10^{-8}$ | 0.45 |
| rs2736990 | PD | $1.36 \times 10^{-7}$ | 1.73 |
| rs393152 | PD | $4.75 \times 10^{-8}$ | 4.68 |

**Table 10:** New loci found by the WTCCC2 genome-wide association studies. We provide the SNP rsid, the disease with which it was associated, the $p$-value reported, and the mean ABF across our analyses. Note that some SNPs appear more than once due to their association with multiple traits. As before, SNPs for which there was summary statistic data for only one trait could not have a MetABF analysis performed, and have in NA the "mean ABF" column.

| SNP | Disease | $p$-value in study | mean ABF |
|---|---|---|---|
| rs17716942 | PS | $4.05 \times 10^{-8}$ | 9.10 |
| rs27524 | PS | $6.81 \times 10^{-10}$ | 14.88 |
| rs240993 | PS | $8.71 \times 10^{-13}$ | 5.79 |
| rs458017 | PS | $8.11 \times 10^{-11}$ | 5.52 |

(Continued on next page)

*(continued from previous page)*

| **SNP** | **Disease** | ***p*-value in study** | **mean ABF ($\log_{10}$)** |
|---|---|---|---|
| rs11581062 | MS | $3.7 \times 10^{-10}$ | 4.62 |
| rs11129295 | MS | $2.3 \times 10^{-8}$ | 4.93 |
| rs669607 | MS | $2.9 \times 10^{-11}$ | 5.99 |
| rs9282641 | MS | $1.5 \times 10^{-9}$ | 6.04 |
| rs11154801 | MS | $1.5 \times 10^{-12}$ | 5.21 |
| rs17066096 | MS | $3.4 \times 10^{-10}$ | 4.00 |
| rs1738074 | MS | $5.3 \times 10^{-11}$ | 6.28 |
| rs4902647 | MS | $3.8 \times 10^{-8}$ | 6.64 |
| rs2119704 | MS | $3.5 \times 10^{-10}$ | 6.04 |
| rs1077667 | MS | $2.1 \times 10^{-12}$ | 6.40 |
| rs874628 | MS | $4.3 \times 10^{-8}$ | 2.02 |
| rs601342 | UC | $3.2 \times 10^{-13}$ | NA |
| rs188755755 | BS pneumo. | $8.91 \times 10^{-9}$ | NA |
| rs140817150 | BS pneumo. | $7.25 \times 10^{-9}$ | NA |
| rs116432683 | BS pneumo. | $7.2 \times 10^{-9}$ | NA |
| rs9869826 | BS overall | $2.88 \times 10^{-8}$ | 1.74 |
| rs9879725 | BS overall | $2.36 \times 10^{-8}$ | -1.12 |
| rs1884318 | BS overall | $4.35 \times 10^{-8}$ | NA |
| rs4632900 | BS overall | $2.96 \times 10^{-8}$ | -0.52 |

*(Continued on next page)*

(continued from previous page)

| **SNP** | **Disease** | ***p*-value in study** | **mean ABF ($\log_{10}$)** |
|---|---|---|---|
| rs75698727 | BS overall | $2.92 \times 10^{-8}$ | NA |
| rs113892119 | BS overall | $5.08 \times 10^{-13}$ | NA |
| rs12295158 | BS overall | $1.32 \times 10^{-10}$ | NA |
| rs334 | BS overall | $1.33 \times 10^{-10}$ | NA |
| rs9271252 | VL India | $1.26 \times 10^{-9}$ | NA |
| rs9271255 | VL India | $1.71 \times 10^{-9}$ | NA |
| rs9271842 | VL India | $1.49 \times 10^{-10}$ | NA |
| rs9271858 | VL India | $1.57 \times 10^{-10}$ | NA |
| rs9272070 | VL India | $1.66 \times 10^{-10}$ | NA |
| rs9271252 | VL Brazil | $2.33 \times 10^{-5}$ | NA |
| rs9271255 | VL Brazil | $2.36 \times 10^{-5}$ | NA |
| rs9271842 | VL Brazil | 0.000194 | NA |
| rs9271858 | VL Brazil | 0.000201 | NA |
| rs9272070 | VL Brazil | $3.68 \times 10^{-5}$ | NA |
| rs356220 | PD | $5.18 \times 10^{-9}$ | 3.52 |
| rs10447854 | PD | $3.11 \times 10^{-9}$ | 3.04 |
| rs417968 | PD | $1.81 \times 10^{-8}$ | 3.90 |
| rs7215239 | PD | $1.49 \times 10^{-8}$ | 4.45 |

#### 6.8 Results of the top regional WTCCC2 SNPs

| SNP | Chr | Position | log10 Mean ABF | Corr. Mat | σ |
|---|---|---|---|---|---|
| rs3890745 | 1 | 2543484 | 8.258 | 3 | 0.1 |
| rs6670226 | 1 | 20067289 | 5.097 | 1 | 0.2 |
| rs10903122 | 1 | 25176163 | 5.137 | 7 | 0.1 |
| rs11209026 | 1 | 67478546 | 14.512 | 2 | 0.4 |
| rs12746893 | 1 | 92983464 | 6.236 | 2 | 0.2 |
| rs12746227 | 1 | 101044667 | 5.361 | 2 | 0.2 |
| rs1335532 | 1 | 116902480 | 6.227 | 7 | 0.1 |
| rs12128408 | 1 | 152755157 | 4.119 | 1 | 0.1 |
| rs7522462 | 1 | 199148218 | 8.425 | 2 | 0.1 |
| rs4672507 | 2 | 62424077 | 16.516 | 1 | 0.4 |
| rs17716942 | 2 | 162968937 | 9.096 | 7 | 0.2 |
| rs11129295 | 3 | 27763784 | 4.928 | 2 | 0.1 |
| rs669607 | 3 | 28046448 | 5.991 | 9 | 0.1 |
| rs11715915 | 3 | 49430334 | 4.277 | 3 | 0.2 |
| rs2173623 | 3 | 55571956 | 4.014 | 4 | 0.1 |
| rs1398607 | 3 | 103238428 | 4.900 | 2 | 0.1 |
| rs7650774 | 3 | 120687740 | 4.023 | 7 | 0.1 |
| rs1920298 | 3 | 123080578 | 4.077 | 12 | 0.1 |
| rs9282641 | 3 | 123279458 | 6.036 | 2 | 0.2 |
| rs860413 | 5 | 35978799 | 4.959 | 2 | 0.1 |
| rs17234657 | 5 | 40437266 | 10.076 | 3 | 0.1 |
| rs30187 | 5 | 96150086 | 17.263 | 2 | 0.2 |
| rs10059288 | 5 | 158791476 | 14.523 | 1 | 0.2 |
| rs1177441 | 6 | 25967392 | 7.354 | 1 | 0.4 |
| rs3173197 | 6 | 90713028 | 4.552 | 12 | 0.2 |
| rs240993 | 6 | 111780407 | 5.786 | 4 | 0.1 |
| rs11154801 | 6 | 135781048 | 5.206 | 2 | 0.1 |
| rs17066096 | 6 | 137494601 | 4.001 | 7 | 0.1 |
| rs7746779 | 6 | 138196194 | 4.999 | 7 | 0.1 |
| rs1738074 | 6 | 159385965 | 6.278 | 7 | 0.1 |
| rs4730280 | 7 | 107337182 | 4.427 | 4 | 0.1 |
| rs4410871 | 8 | 128884211 | 4.254 | 12 | 0.1 |
| rs3812558 | 9 | 138388956 | 4.246 | 2 | 0.2 |
| rs4147359 | 10 | 6148445 | 4.543 | 2 | 0.1 |
| rs793108 | 10 | 31455112 | 4.768 | 4 | 0.1 |
| rs1250552 | 10 | 80728033 | 4.054 | 7 | 0.1 |
| rs10837587 | 11 | 41132173 | 4.163 | 3 | 0.4 |
| rs16920014 | 11 | 93566105 | 13.683 | 3 | 0.4 |
| rs1800693 | 12 | 6310270 | 8.332 | 7 | 0.1 |
| rs4902647 | 14 | 68323944 | 6.636 | 12 | 0.1 |
| rs1372363 | 14 | 87542931 | 6.090 | 2 | 0.2 |
| rs1035089 | 16 | 10955851 | 5.778 | 2 | 0.1 |
| rs2041670 | 16 | 11082153 | 7.031 | 2 | 0.2 |
| rs12445650 | 16 | 30952398 | 4.383 | 12 | 0.1 |
| rs7404688 | 16 | 85390918 | 4.086 | 3 | 0.2 |
| rs744166 | 17 | 37767727 | 4.705 | 7 | 0.1 |
| rs393152 | 17 | 41074926 | 4.685 | 1 | 0.2 |
| rs17769552 | 17 | 41283070 | 5.508 | 1 | 0.2 |
| rs1981997 | 17 | 41412603 | 4.675 | 1 | 0.2 |
| rs2668692 | 17 | 41648797 | 4.245 | 1 | 0.2 |
| rs1077667 | 19 | 6619972 | 6.397 | 2 | 0.2 |
| rs8106664 | 19 | 10589030 | 4.566 | 12 | 0.1 |
| rs676388 | 19 | 53903781 | 4.410 | 7 | 0.1 |
| rs6138362 | 20 | 24684866 | 4.988 | 1 | 0.2 |
| rs2094871 | 21 | 39382729 | 15.511 | 12 | 0.2 |

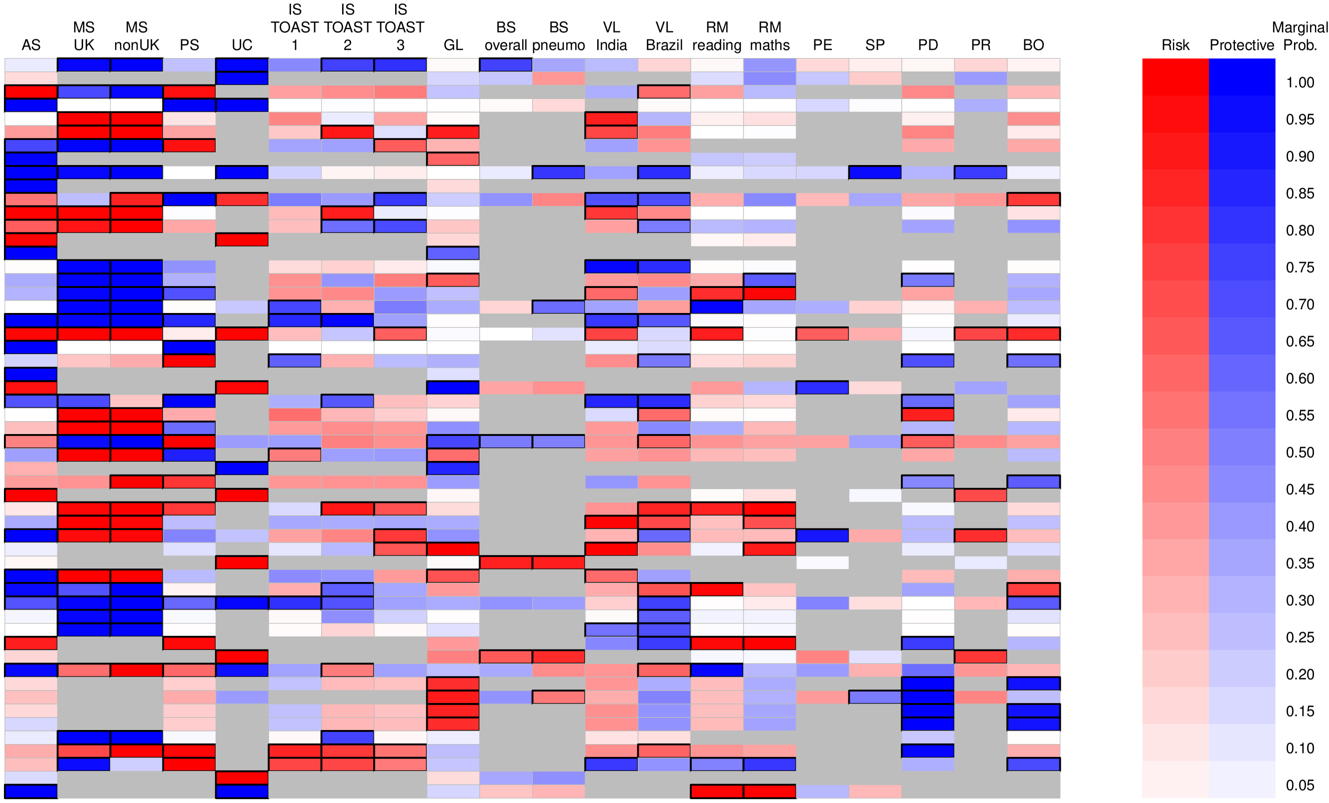


**Figure 15:** Table showing the SNPs that we took forward for further analysis of their cross-phenotype associations. In the table on the left, we give the rsid, chromosome, genomic position (on NCBI Build 36), the $\log_{10}$(mean ABF), the correlation matrix and value of $\sigma$ associated with the highest ABF across all possible subsets and calculated using every prior correlation matrix and value for $\sigma$. In the chart on the right, we show the pattern of association that corresponded to the highest Bayes factor using the subset exhaustive approach and the prior parameters in the table at the left. Black boxes are drawn around the traits associated with the SNP and the color indicates whether the effect allele was a risk (red) or protective allele (blue). The intensity of the color is based on the marginal probability of association at the SNP, with the scale at the right.

Marginal probabilities of association were calculated by calculating the ABF for every possible subset of association using the priors associated with the highest ABF overall. These ABFs were normalized to get a probability distribution over all the models (i.e. there was a flat prior on the models of association). Marginal probabilities of association for each trait were calculated using this distribution.

#### 6.9 Colocalised signals

Our results do not differentiate between a single variant with multiple true associations, and true associations at many individual SNPs in LD with one another, which all colocalise at a single SNP. For example, in Figure 16, the SNP rs8106664 (vertical black line) had the highest ABF in the region. However, the strongest associations, which are in the MS non-UK and PS cohorts, actually lie upstream of the SNP. Additionally, there are possible associations with the IS cohorts lying both upstream and downstream of the SNP. It therefore seems plausible that ABF at this SNP is the result of colocalisation of markers in LD with each other at rs8106664, rather than association of each phenotype with this particular SNP.

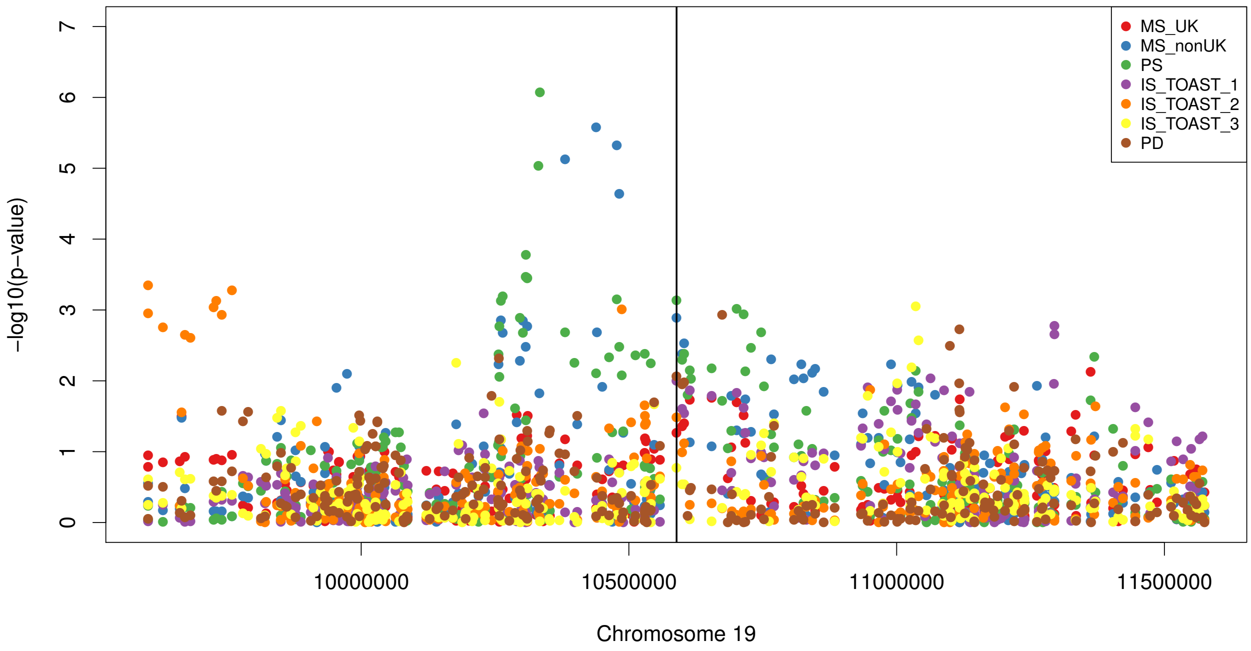


**Figure 16:** Plot of $-\log_{10}(p\text{-values})$ for the seven WTCCC2 traits which show evidence of association with rs8106664 (black vertical line) on chromosome 19.

### 7 Comparisons to CPBayes and MTAG

Using simulated data, we compared our subset exhaustive approach to two other methods, CPBayes (Majumdar, Haldar, Bhattacharya, & Witte, 2018) and MTAG (Turley et al., 2018). Our simulated datasets contained 60,000 independent SNPs across five studies. The first 10,000 SNPs were simulated to have no true effects in any of the studies, the next 10,000 SNPs were simulated to have true effects in the first study, the next 10,000 SNPs were simulated to have true effects in the first and second studies, and so on so that the last 10,000 SNPs were simulated to have true effects in all studies.

Each study was assumed to have 3000 cases and 6000 controls. We investigated the case where each study was completely independent of the others (i.e. no samples shared across studies) and the case where each study shared 3000 controls with each of the others. True effect sizes were generated from a multivariate normal distribution centered on the zero vector. True effect sizes were simulated to have uniform correlation coefficients between each study, whose values were either 0 or 0.5. Similarly, the standard error of true effect sizes was uniformly 0.2 or 0.4, and the effect allele frequency was uniformly 0.1 or 0.5. We used two different covariance matrices for all analyses: one where the cryptic correlation between studies had been calculated analytically, using all markers whose $p$-values were greater than 0.1, and one where the correlation between studies was calculated based on the number of shared controls using the function provided by CPBayes. Because only one sixth of the SNPs in our simulated dataset were truly null, we would expect the correlations calculated based on the number of shared controls to be the more accurate of the two methods.

We attempted to create datasets where true effects were uniformly 1.05 or 1.1. However, we did not include the results in our analysis, as the resulting distribution of effect size estimates had a median value significantly higher than 0, causing the MTAG analysis to fail. We additionally do not report the results of the uncorrelated CPBayes analysis, as these were similar to those obtained from using the correlated analysis with a diagonal correlation matrix.

In total, there were 32 analyses of 16 datasets for CPBayes and MTAG. Using our method, we analyzed the data using priors values of $\rho = 0$ and $\sigma = 0, 2$, which assume independent effects of moderate size across all studies. We performed the subset exhaustive analysis using a flat prior on the models of association and then normalized the results at each simulated SNP to get the distribution of posterior probabilities. From this distribution, the marginal probability of association for each trait at each simulated SNP was calculated.

We investigated the inferences made about the marginal evidence of association for a given trait at a given marker. For both CPBayes and MetABF, these are marginal probabilities of association, while MTAG produces $p$-values of association. We calculated the appropriate thresholds for a given false positive rate for each method analytically and then plotted the corresponding true positive association rates in Figure 17. We also plot the run time of each method (wall time) in Figure 18.

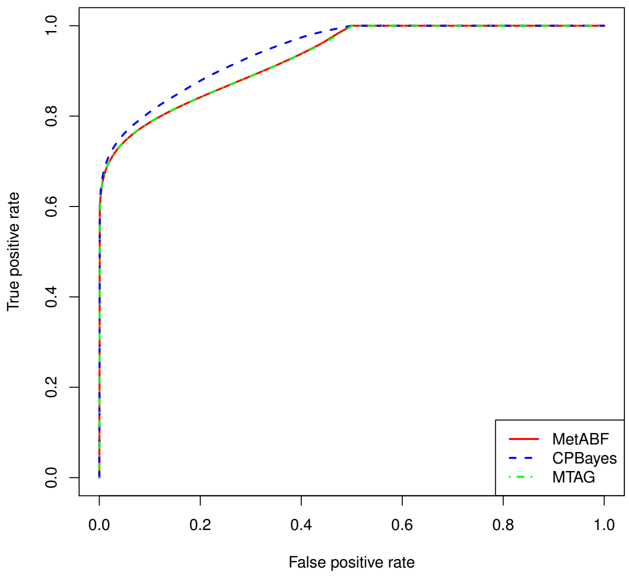


**Figure 17:** Comparison of the ability of each method (colored lines) to detect true positives ($y$-axis) for a given false positive ($x$-axis) rate. False positive rates for each method were determined analytically for each method for the association statistic between a given trait and a given marker. For CPBayes and MetABF, this was the marginal probability of association and for MTAG, this was the $p$-value of association.

#### 7.1 CPBayes

As we see in Figure 17, CPBayes has a slightly higher true positive rate than the other two methods for all false positive rates less than 0.5. However, it also had the longest run time of the three methods, taking between 3 and 12 days to analyze a dataset of 60,000 markers, instead of the seconds or minutes required by the other two methods. When directly comparing the most likely models of association according to our approach with the

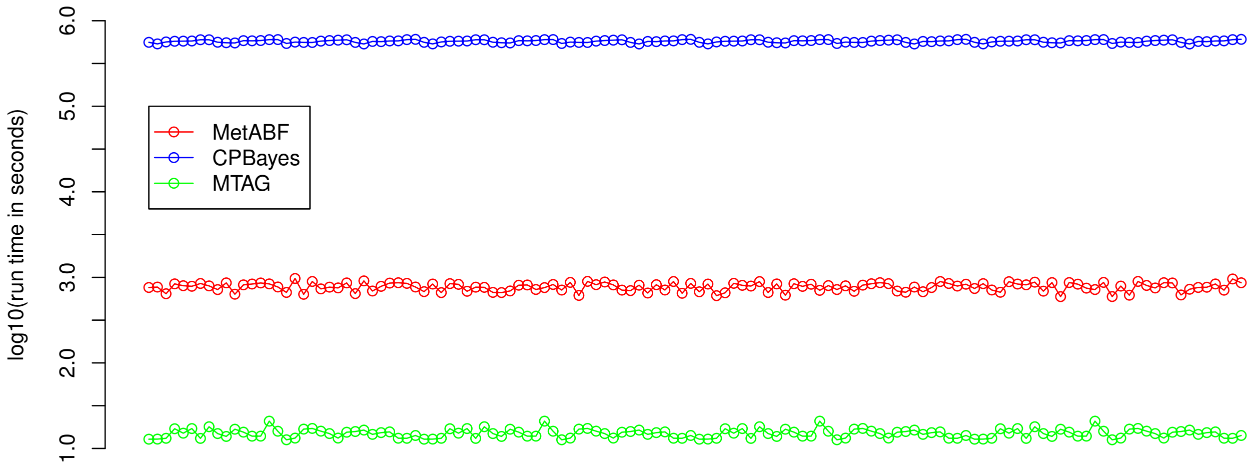


**Figure 18:** Plot comparing the run times in $\log_{10}(seconds)$ ($y$-axis) for each analysis (colored lines) on each dataset ($x$-axis). Here, 1 corresponds to 10 seconds, 3 to 16 minutes and 40 seconds, 5.5 to 3 days, 15 hours, 50 minutes, and 27.8 seconds, while 6 corresponds to 11 days, 13 hours, 46 minutes, and 40 seconds.

one produced by CPBayes, these two were often in agreement.

MetABF's subset-exhaustive approach allows us to explore the entire model space. This means that in addition to calculating posterior probabilities of association, we can also use the intersection of the top $k$ models of association or take the $Q\%$ credible set. MetABF approach takes an average of just under 8 minutes to analyze the full 60,000 markers, while CPBayes takes an average of 2.3 days to run when it assumes no cryptic relatedness between any of the studies in the meta-analysis (not shown in our plots), and 6.6 days to run when the method uses a cryptic correlation matrix. While both methods can easily be parallelized—since each one analyzes each SNP in isolation—it is difficult to see how CPBayes could scale easily to genome-wide data. It is also not clear that the performance of this method compared to that of the other two is so much better as to be worth the increased run time.

#### 7.2 MTAG

We note that MTAG and MetABF perform very similarly at a given false positive rate, with the two methods generally differing from one another by no more than 0.005. In terms of run time, MTAG is easily the fastest of the three methods (Figure 18), taking an average of just over 15 seconds to run each of its analyses. Had MTAG been able to calculate the genetic covariance due to shared controls, this might have added another 5-10 seconds to the run time, which would still have made it faster than the other methods. However, these figures do not take into account the amount of time that it takes to create the required reference panels and to calculate LD scores. While the software and the makers of LD score regression (B. Bulik-Sullivan et al., 2015; B. K. Bulik-Sullivan et al., 2015) do provide reference panels, our struggle to generate appropriate simulations of these extra data reflect the difficulties one might encounter if a marker of interest is not on one of the provided reference panels, or if one or more of the studies in question were conducted in a population for which there is no good reference panel.

Calculation of LD scores requires genotype data that may not be available to a researcher who has access only to GWAS summary statistics. It is also not clear what reference panel is appropriate for meta-analyses across different ethnic groups. Additionally, due to the close connection between MTAG and LD score regression, this method assumes that the summary statistic data available will contain at least 200,000 SNPs across the genome and that there will be some correlation structure among these markers. Our simulated datasets violated these assumptions, and while we were able to create dummy LD score regression reference panels, safe in the knowledge that there was no true underlying correlation among our markers, it is unclear what someone with real data ought to do when dealing with fewer than 200,000 markers, trans-ethnic data, or when there is no genetic data and the markers of interest are not represented on a pre-calculated reference panel. This need to take advantage of the underlying correlation structure also means that MTAG, unlike the other two methods, cannot be used to perform analyses on subsets of SNPs, and in particular, cannot be used on traits or SNP subsets where the median signed test statistic is significantly higher than zero.

### Figure Legends

noindent Figure 14:

Figure 15:

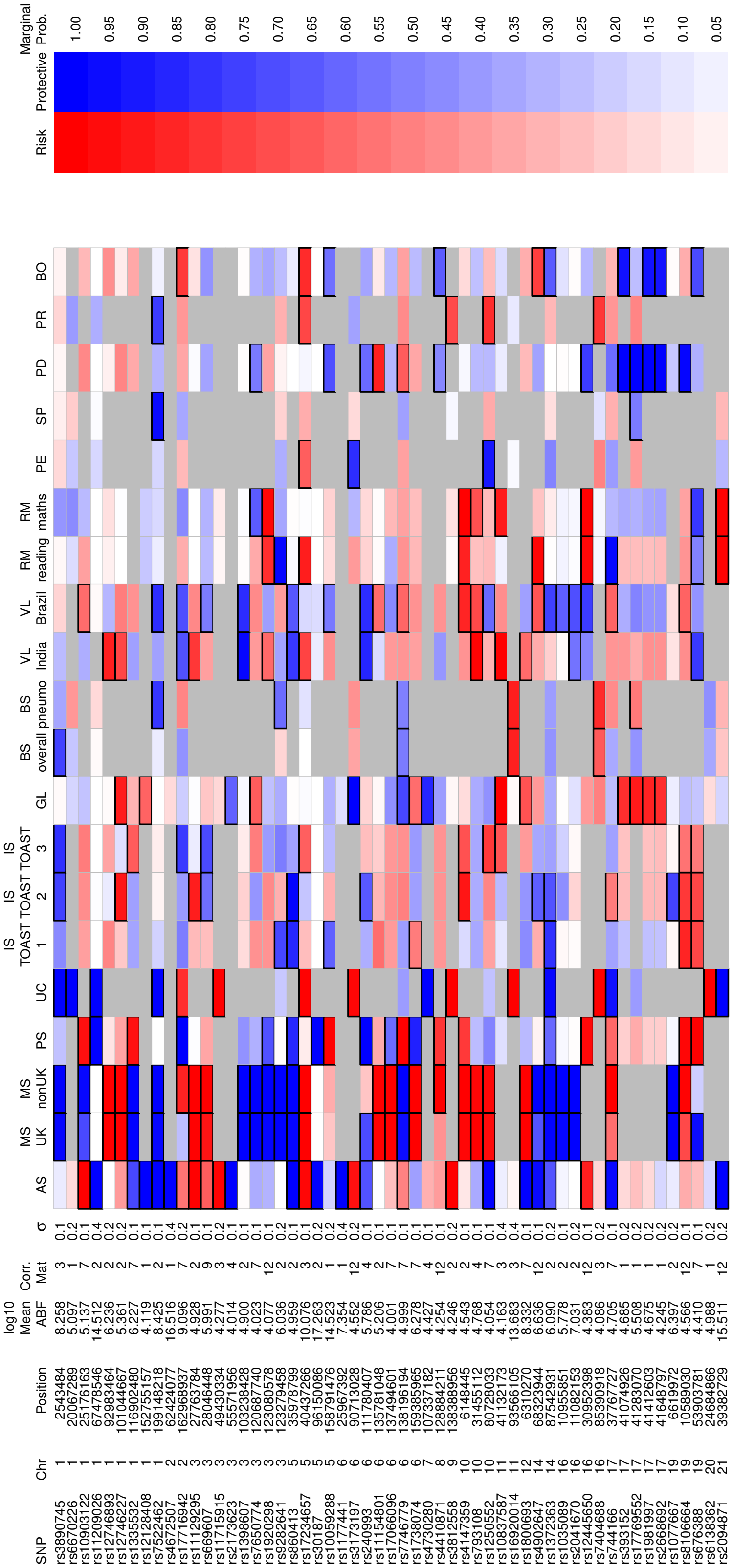

Marginal Prob.
Protective
Risk
1.00
0.95
0.90
0.85
0.80
0.75
0.70
0.65
0.60
0.55
0.50
0.45
0.40
0.35
0.30
0.25
0.20
0.15
0.10
0.05
BO
PR
PD
SP
PE
RM maths
RM reading
VL Brazil
VL India
BS pneumo
BS overall
GL
IS TOAST 3
IS TOAST 2
IS TOAST 1
UC
PS
MS nonUK
MS UK
AS
σ
Corr. Mat
log10 Mean ABF
Position
Chr
SNP

Figure 16:

Figure 17:

Figure 18:
